# RBBP4 modulates gene activity through acetylation and methylation of histone H3 lysine 27

**DOI:** 10.1101/2021.09.09.459568

**Authors:** Weipeng Mu, Noel S Murcia, Keriayn N. Smith, Debashish U Menon, Della Yee, Terry Magnuson

**Affiliations:** Department of Genetics, and Lineberger Comprehensive Cancer Center, The University of North Carolina at Chapel Hill, Chapel Hill, North Carolina, NC 27599-7264, USA

**Keywords:** histone methylation, histone acetylation, transcriptional regulation, Polycomb-Group proteins, enhancers, super-enhancer, epigenetic memory

## Abstract

RBBP4 is a core subunit of polycomb repressive complex 2 (PRC2) and HDAC1/2-containing complexes, which are responsible for histone H3 lysine 27 (H3K27) methylation and deacetylation respectively. However, the mechanisms by which RBBP4 modulates the functions of these complexes remain largely unknown. We generated viable mouse embryonic stem cell lines with RBBP4 mutations that disturbed methylation and acetylation of H3K27 on target chromatin and found that RBBP4 is required for PRC2 assembly and H3K27me3 establishment on target chromatin. Moreover, in the absence of EED and SUZ12, RBBP4 maintained chromatin binding on PRC2 loci, suggesting that the pre-existence of RBBP4 on nucleosomes serves to recruit PRC2 to restore H3K27me3 on newly synthesized histones. As such, disruption of RBBP4 function led to dramatic changes in transcriptional profiles. In spite of the PRC2 association, we found that transcriptional changes were more closely tied to the deregulation of H3K27ac rather than H3K27me3 where increased levels of H3K27ac were found on numerous cis-regulatory elements, especially putative enhancers. These data suggest that RBBP4 controls acetylation levels by adjusting the activity of HDAC complexes. As histone methylation and acetylation have been implicated in cancer and neural disease, RBBP4 could serve as a potential target for disease treatment.

## Introduction

Histone modifications are key epigenetic regulators that specify the transcriptome during lineage determination and tightly control transcriptional states to preserve cellular identity over cell generations (1). Of the various types of histone modifications, methylation and acetylation are central regulators of gene expression. These modifications are functionally deterministic on chromatin regions around cis-regulatory elements such as promoters, enhancers, and silencers, where they impact transcription factor binding and activity by altering chromatin structure and accessibility (2, 3). Lysine 27 on histone H3 (H3K27) can be either methylated by Polycomb repressive complex 2 (PRC2) (4) or acetylated by p300 and CBP (5), and these modifications serve as hallmarks of transcription repression and activation respectively. Methylation and acetylation of H3K27 can also occur on nucleosomes marked by other histone modifications, such as lysine 4 methylation of histone H3 (H3K4me) that regulate similar or differing effects on transcription. These data suggest that crosstalk between histone modifications regulates chromatin function (6). Disruption of H3K27 methylation and acetylation results in numerous cellular defects, including cell proliferation and differentiation anomalies during embryogenesis and tissue specification (7–9). Their dysregulation has also been implicated in disease processes such as tumorigenesis (10).

PRC2, the methyltransferase with specific activity toward H3K27 is responsible for the mono-, di- and tri-methylation modifications of this residue (H3K27me1/me2/me3) (4). PRC2 maintains gene silencing primarily through the deposition of H3K27me3 at promoter regions (11, 12). This activity has significant developmental implications. For example, studies have shown that the H3K27me3 accumulated on key developmental genes are not replaced by protamines in sperm (13, 14) and can be transmitted to the next generation to regulate embryonic gene expression (15). Since mammals do not have polycomb repressive elements (PREs) with specific cis-regulatory sequences for polycomb group (PcG) binding, how PRC2 is bound to specific sites in the genome for the establishment of H3K27me3 during mammalian development remains an open question.

Newly replicated chromatin echoes the parental histone modification landscape with modified parental histones being efficiently and accurately recycled to sister chromatids during chromatin replication (16). This restoration of histone modifications during cell proliferation plays a vital role in the maintenance of cellular identity and homeostasis. Parental H3K27me3 modifications are re-incorporated in proximity to their original positions during DNA replication, suggesting that existing marks are docking sites for PRC2 recruitment to its target loci during cell division (17). Surprisingly however, recent studies have demonstrated that the preexistence of parental H3K27me3 is not required for propagation of this histone mark during cell proliferation (18). Although ancillary PRC2 subunits, including JARID2, AEBP2, PCL1-3, EPOP, and PALI, are also implicated in PRC2 recruitment, depletion of these subunits caused only subtle alterations in the enrichment of H3K27me3 on target genes (19). These findings led us to speculate that certain nucleosome-associated proteins may guide PRC2 to specific genomic sites.

RBBP4 and RBBP7 are core subunits of PRC2 that share 92% identity. Both proteins contain WD40 domains that serve as scaffolds for protein complex assembly or platforms to recruit diverse molecules that form functional complexes (20, 21). As histone chaperones, RRBBP4/7 directly bind to histones H3 and H4 and are proposed to act as escorting factors that connect protein complexes to their substrate nucleosomes. They have also been implicated in multiple aspects of the cellular events that are involved in histone metabolizing processes depending on individual complex composition (20, 22). Within PRC2, RBBP4 physically interacts with SUZ12 and AEBP2 (23). However, unlike EED, SUZ12 and EZH2, the RBBP4/7 homologs are not required for methyltransferase activity (24), and precise roles for RBBP4/7 in regulating PRC2 functions remain unclear. One key question is whether RBBP4/7 determine site-specific recruitment of PRC2 to chromatin for the establishment and maintenance of H3K27 methylation during cell division.

RBBP4/7 are also components of histone deacetylase (HDAC) complexes that are essential for normal development, including NuRD, Sin3, and CoREST (25). RBBP4/7, along with HDAC1 and HDAC2, are shared among these complexes and coordinate with other complex-specific subunits to provide target specificity or additional catalytic activities (10). In accordance with the association between histone acetylation and transcriptional activation, HDAC complexes erase histone acetylation marks to function as corepressors that transcriptionally repress target genes. It has been shown that dynamic acetylation and deacetylation are required for active transcription to occur, indicating that HDAC-complexes also function in transcriptional activation (25, 26). The NuRD complex, which is comprised of a dimer of the sub-components HDAC1:RBBP4:MTA1, binds nucleosomes, with the RBBP4 protein mediating interaction with histone H3 tails as a mechanism for recruitment of NuRD to chromatin (27). Studies to understand how RBBP4 interacts within different HDAC complexes to regulate histone acetylation genome-wide are needed to clarify the regulation of histone acetylation and deacetylation in transcription. Also, since RBBP4/7 are core components of PRC2 subcomplexes and HDAC1/HDAC2-containing complexes, their molecular and functional characterization can help us to understand differential binding-site specificity, and PRC2/HDAC recruitment to chromatin.

Since RBBP4/7 are essential for cell viability, it is difficult to test their roles experimentally. By utilizing CRISPR-Cas9 to target RBBP4/7 in mouse embryonic stem cells (ESCs), we generated viable ESC colonies with mutated RBBP4/7. We found that RBBP4 mutations remarkably altered transcriptional profiles and the landscape of H3K27 methylation and acetylation. Our results suggest that RBBP4 marks PRC2 target chromatin for site-specific recruitment of other PRC2 subunits for the trimethylation of H3K27. RBBP4 also modulates the catalytic activity of HDAC-containing complexes and controls H3K27ac levels on cis-regulatory elements, especially enhancers to finely tune gene activity. Altogether, our findings demonstrate that RBBP4 plays broad and differential roles in the regulation of functionally different chromatin modifying complexes.

## Results

### 1. RBBP4 is required for efficient PRC2 binding and H3K27 trimethylation on target loci

To generate viable and proliferating cell lines with disrupted RBBP4 or RBBP7 function, we utilized the CRISPR-Cas9 technique to target the N-terminus of RBBP4 and RBBP7 with sgRNAs in mouse ESCs. Mutant cell lines were validated by examining nucleotide sequences and protein expression levels of *Rbbp4* and *Rbbp7*. For *Rbbp4*, we obtained two ESC clones; the first mutant named *Rbbp4^Ins/Ins^,* carries a missense mutation (Val84Asp) and a Cys insertion at the 85^th^ amino acid (Figure S1A), and the second mutant named *Rbbp4^Ins/Del^*, has *Rbbp4^Ins^* on one allele and a six amino acid deletion (79^th^ to 84^th^) on the other allele (Figure S1B). Western blot analysis indicated that mutant RBBP4 proteins were expressed at lower levels and RBBP4^Ins^ migrated slower relative to controls (Figure 1A). The gel-shift associated with RBBP4^Ins^ was possibly caused by the oxidation of the inserted cysteine because cysteine oxidation can retard protein migration through SDS-PAGE (28). We did not retrieve *Rbbp4* homozygous knockout clones, confirming *Rbbp4* as an essential gene (24). For *Rbbp7*, we obtained two homozygous knockout ESC clones with no RBBP7 protein detected (Figure 1B), suggesting RBBP7 is not required for ESC viability. The mutations in *Rbbp4* or loss of RBBP7 did not impair the expression of other PRC2 subunits and global levels of H3K27 methylation as assessed by Western blot (Figure 1A-C).

**Figure 1.**
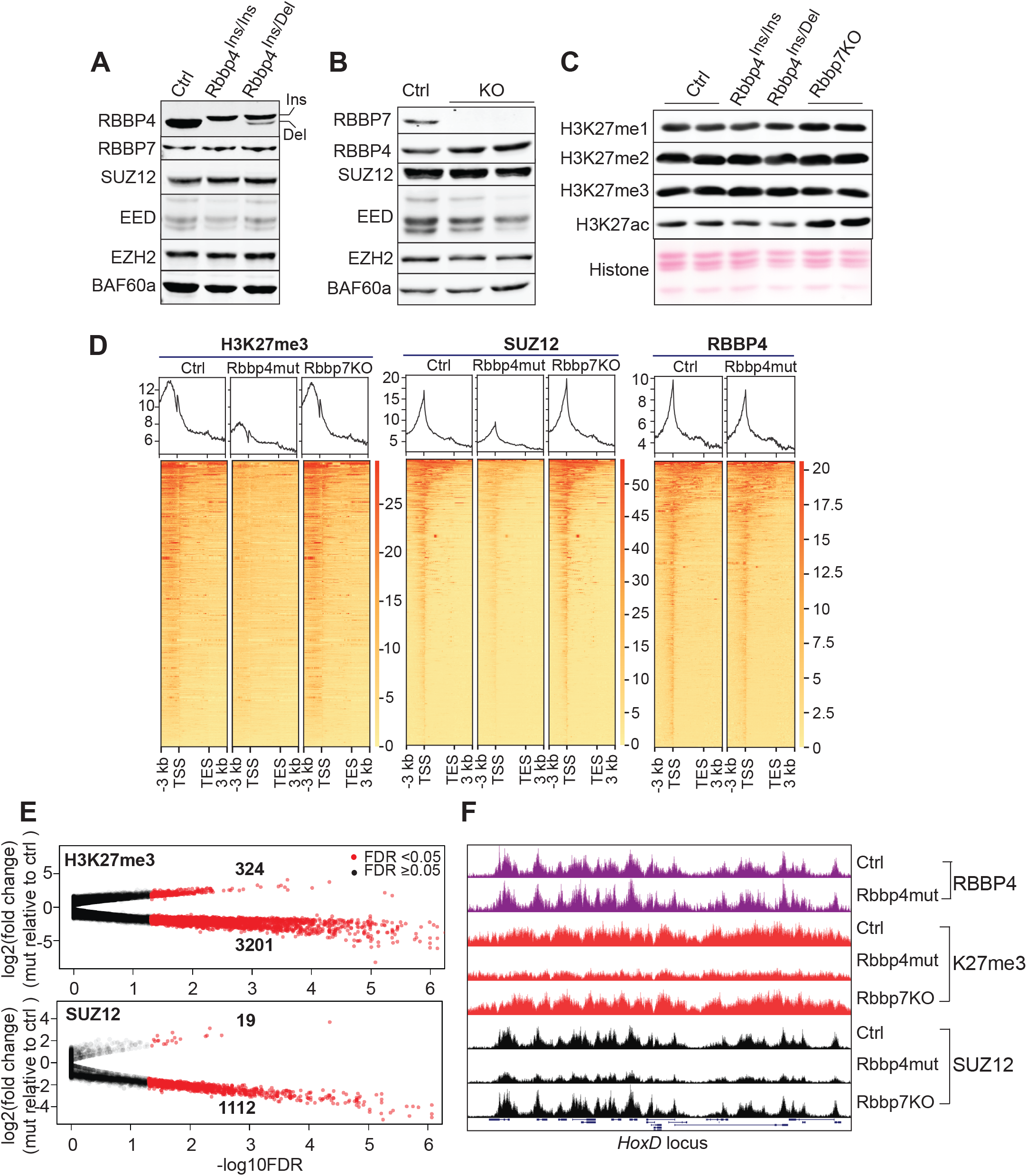
RBBP4 is required for SUZ12 binding and trimethylation of H3K27 on PRC2 target loci. (A) Western blot analysis of RBBP4 and other PRC2 core subunits in E14 (ctrl) and Rbbp4 mutants (mut) ESCs. BAF60a serves as a loading control. (B) Western blot analysis of RBBP7 and other PRC2 core subunits in E14 (ctrl) and Rbbp7 knockout (KO) ESCs. (C) Western blot analysis of H3K27 methylation and acetylation in E14 (ctrl), Rbbp4 mutant (mut), and Rbbp7 knockout (KO) ESCs. (D) Profile plots (top) and heat maps (bottom) depicting H3K27me3, SUZ12 and RBBP4 enrichment across gene bodies. Genes are ranked based on the read density, and Rbbp4mut and Rbbp7ko use the same gene order as the controls (ctrl). TSS - transcription start site, TES – transcription end site. (E) Differential analysis of H3K27me3 and SUZ12 enrichment in controls and Rbbp4 mutants. Each dot represents a genomic region with H3K27me3 or SUZ12 binding. (F) Genome browser images of HoxD locus marked by H3K27me3, RBBP4, and SUZ12.

We performed ChIP analyses to examine H3K27me3 enrichment and PRC2 binding on PRC2 target loci. Unlike global H3K27me3 levels, the enrichment of H3K27me3 across gene bodies was dramatically reduced in *Rbbp4* mutants compared to the controls (Figure 1D). This was accompanied by a decrease in SUZ12 (Figure 1D) and EZH2 binding (Figure S1C). We confirmed ChIP-seq results by ChIP-qPCR analysis on genes with H3K27me3 localized broadly across gene bodies (*Hoxa10*, *Hoxd9*, *Pax7*, *T*) or narrowly around the TSS (*Runx2*, *Shc3*, *Wnk2*) (Figure S1D, S1E). In contrast, loss of RBBP7 did not affect genomic patterns of H3K27me3 and SUZ12 (Figure 1D). Together these data signify that RBBP4 is essential for PRC2 binding on target loci for H3K27me3.

Differential binding analysis revealed 3201 and 1112 genomic loci with decreased H3K27me3 and SUZ12 respectively (Figure 1E). Despite these differences, the enrichment of mutated RBBP4 was similar to wildtype RBBP4 on PRC2 loci (Figure 1D, 1F), suggesting that mutant RBBP4 remained nucleosome bound. Although global levels of H3K27me3 in mutant cells were similar to control cells, we did not detect redistribution of H3K27me3 and PRC2 on chromatin to compensate for the reduction across gene bodies. For instance, *Fancg* and *Mybl1*, two germ cell specific genes, which are inactive but not marked by PRC2 in ESCs did not show increased H3K27me3 accumulation and SUZ12 binding in *Rbbp4* mutants (Figure S1D, S1E). In mouse ESCs, the major population of loci with depleted H3K27me3 are key developmental genes such as the Hox gene clusters (Figure 1F), which have bivalent domains that also contain active H3K4me2/3 (Figure S1F). Fewer genomic regions were identified with increased SUZ12 and H3K27me3 occupancy in RBBP4 mutants (Figure 1E). These sites displayed narrow SUZ12 and H3K27me3 enrichment that failed to meet the threshold of commonly used peak caller, MACS2 (Figure S2). These results suggest that RBBP4 facilitates target genes H3K27 trimethylation by PRC2.

### 2. Rbbp4 mutations disrupt PRC2 assembly on target chromatin

Recent studies demonstrated that SUZ12 is essential for guiding PRC2 to correct gene targets in the genome, thereby directing the deposition of H3K27 methylation (18, 29). SUZ12 also coordinates with RBBP4 to mediate the formation of PRC2 dimers to enhance binding to CpG islands throughout the genome, and these are hallmark PRC2 binding sites in mammalian cells (23). To test whether decreased SUZ12 on PRC2 targets in *Rbbp4* mutants are related to impaired RBBP4 chromatin binding, we analyzed RBBP4 enrichment specifically on genomic loci with reduced SUZ12 binding. The amount of RBBP4 chromatin binding in *Rbbp4* mutants is equivalent to that of controls (Figure 2A), indicating that the mutations do not disrupt RBBP4 binding to PRC2 targeted nucleosomes. Since the mutations are on the N-terminus of RBBP4, a series of N-terminal truncated RBBP4 constructs were created to map the interaction between RBBP4 and SUZ12. The most N-terminal segment of RBBP4 (1-126 aa) is sufficient to physically interact with PRC2 subunits (Figure 2B), suggesting that RBBP4 mutations on N-terminus interfere with PRC2 assembly on target chromatin.

**Figure 2.**
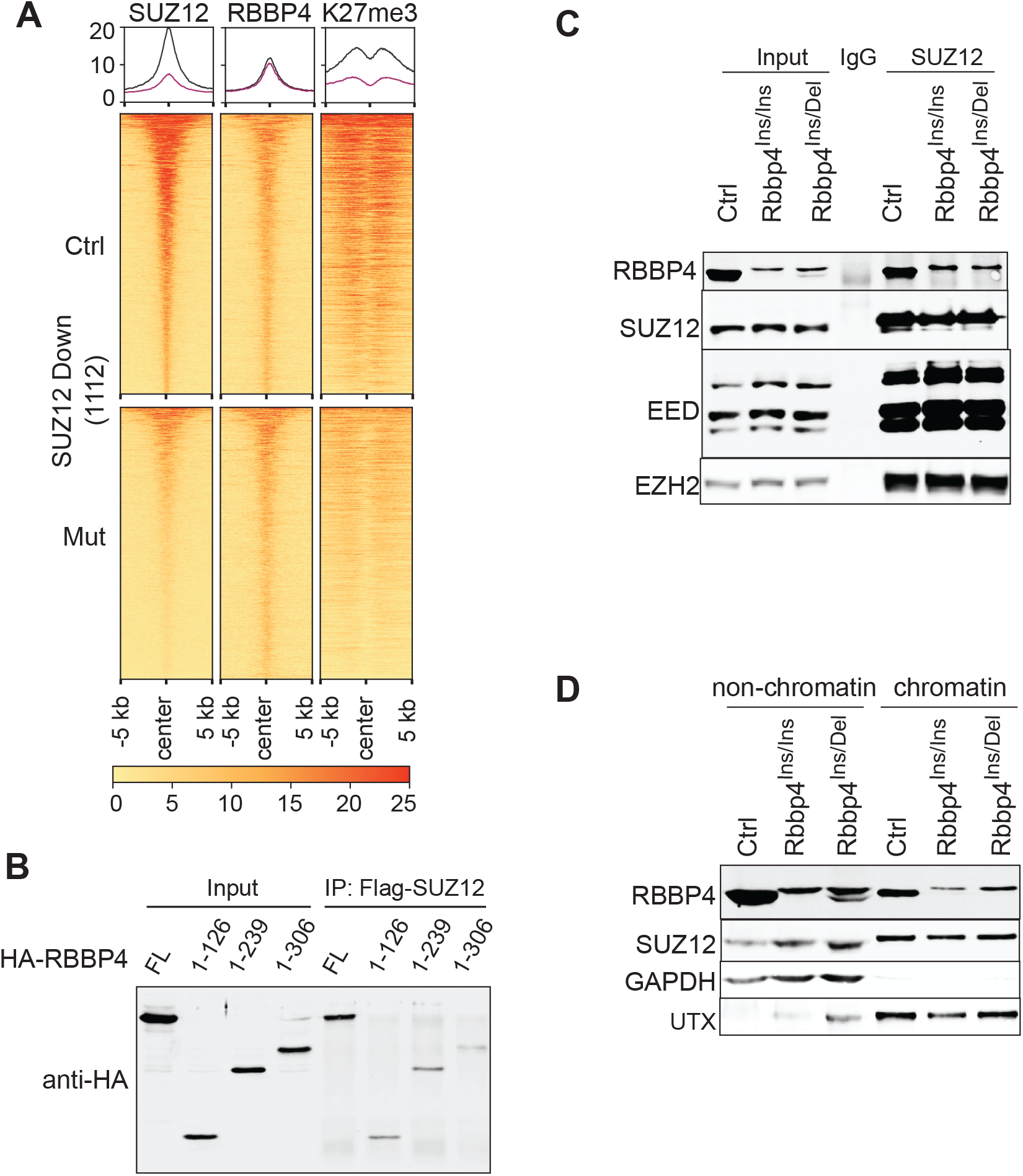
The N-terminus of RBBP4 is essential for recruiting SUZ12 to chromatin. (A) The distribution and enrichment of RBBP4, K27me3, K27ac and HDAC1 within -5/+5 kb of the center of SUZ12-enriched loci. 1112 genomic regions which have decreased SUZ12 binding in the mutants were included for analysis. In the heatmap, the genomic regions are ordered as those in SUZ12. (B) Co-immunoprecipitation and Western blot analysis mapping the interaction betwee N-terminus of RBBP4 and SUZ12. (C) Co-immunoprecipitation and Western blot analysis for assessing the interaction between endogenous RBBP4 mutants and SUZ12. (D) Western blot analysis on cellular fractions of RBBP4 and SUZ12. GAPDH and UTX serve as non-chromatin and chromatin-bound protein controls, respectively.

We performed endogenous co-immunoprecipitation assays using whole cell extracts to determine the effect of mutant RBBP4 on PRC2 subunit incorporation. Although the enrichment of mutant RBBP4 on SUZ12 target loci was similar to that for wild-type RBBP4 (Figure 2A), SUZ12 pulled down less mutant RBBP4 compared to wild-type forms of RBBP4 (Figure 2C). This suggests that RBBP4 mutations interfere with binding of PRC2 subunits to target genomic regions. However, the incorporation of SUZ12, EED and EZH2 into the PRC2 core is not impacted in *Rbbp4* mutant ESCs (Figure 2C), indicating that PRC2 assembly prior to loading onto target loci is independent of RBBP4.

Since dramatic reduction of SUZ12 on PRC2 loci in *Rbbp4* mutants did not accompany with decreased global levels of SUZ12, we tested whether RBBP4 mutations led to the redistribution of SUZ12 in cellular compartments. *Rbbp4* mutant ESCs did not have more SUZ12 in non-chromatin cellular fractions compared to control cells (Figure 2D), suggesting PRC2 subunits are redistributed over chromatin due to RBBP4 disruption. It is possible that SUZ12 is relocated to other genomic regions for H3K27me3. It has been demonstrated that truncated PRC2 that retains catalytic activity but loses binding capability was able to maintain global levels of H3K27me3, and this occurs at lower levels in non-PRC2 target regions and cannot be detected by ChIP-seq (18). These data could explain the patterns of SUZ12 and H3K27me3 in RBBP4 mutant cells. Nevertheless, our findings indicate that RBBP4 plays an important role in assisting PRC2 binding to target chromatin.

### 3. RBBP4 signals recruitment of PRC2 subunits to target loci

It remains unknown whether certain PRC2 subunits mark genomic loci, signaling the assembly of PRC2 and maintenance of H3K27me3 at correct sites during chromatin replication. Although PRC2 ancillary subunits facilitate PRC2 recruitment to target chromatin, maintenance of H3K27me3 on PRC2 target loci does not require their expression (19). EED and SUZ12 are essential for the trimethylation of H3K27. We surmised that loss of H3K27me3 on PRC2 loci could not be recovered if these two subunits were responsible for both marking PRC2 target loci and for recruiting PRC2 to those targets. In order to test this, we generated *Eed* and *Suz12* knockout ESC lines in which H3K27me3 was depleted (Figure 3A). By introducing Flag-tagged EED or SUZ12 into corresponding knockout cell lines, H3K27me3 was restored (Figure 3B) specifically on PRC2 target genes (Figure 3C). In contrast, inactive genes that are not marked by H3K27me3 in ESCs, such as *Fancg* and *Mybl1*, remain deficient in H3K27me3 (Figure 3C). These results suggest that PRC2 target loci are not identified by H3K27me3 chromatin modifications alone and that PRC2 subunits rely on another factor which reads chromatin and guides the organization of PRC2 subunits to their target loci.

**Figure 3.**
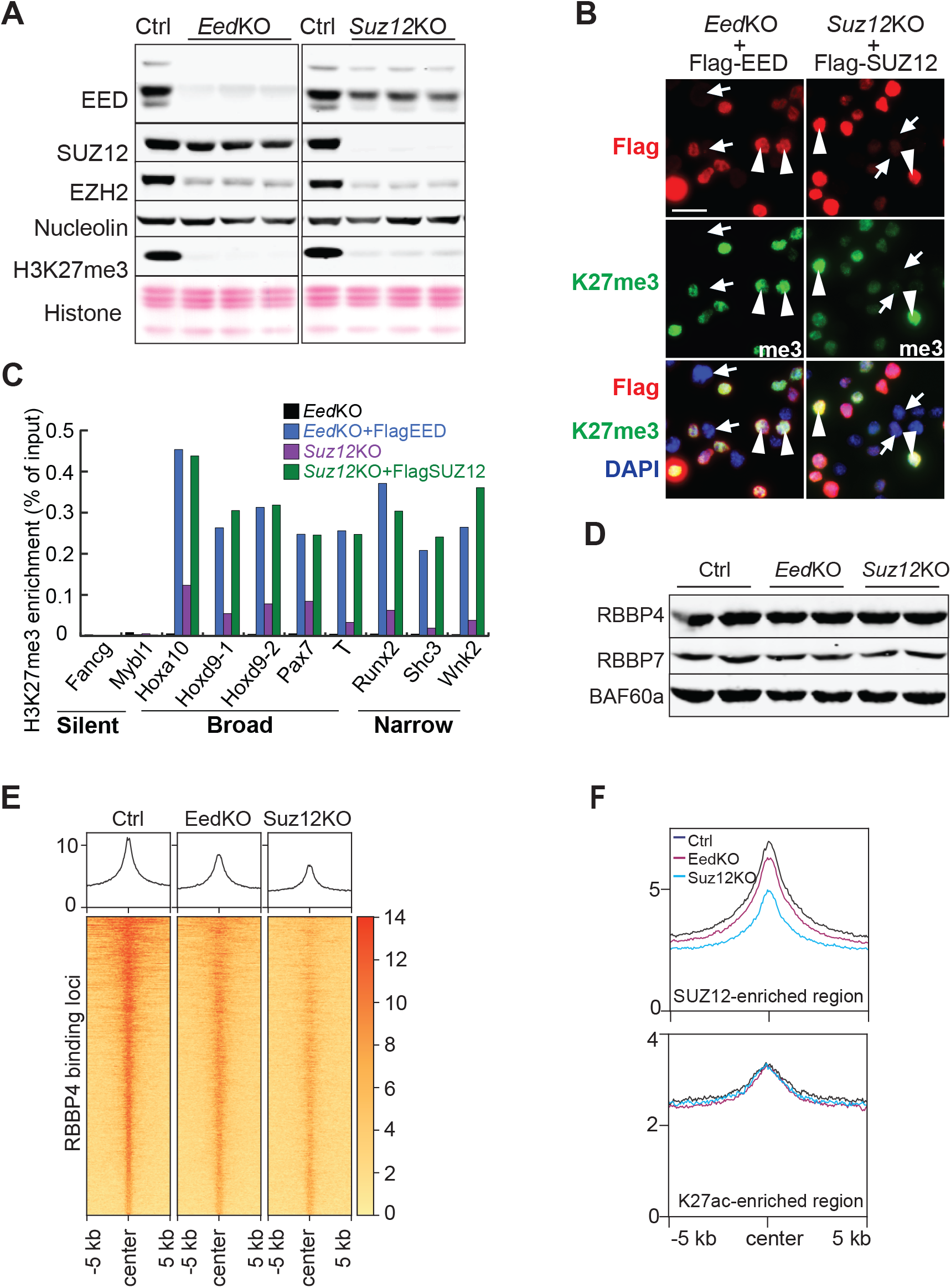
RBBP4 is stable and can associate with PRC2 loci in the absence of SUZ12 and EED. (A) Validation of Suz12 and Eed knockout cell lines by Western blot analysis. Nucleolin serves as nuclear protein control. (B) Immunofluorescence analysis of H3K27me3 in EedKO and Suz12KO ES cells that were transfected with FLAG-tagged EED or SUZ12, respectively. Scale bar: 20 μm (C) Quantitation of H3K27me3 enrichment on PRC2 target genes by ChIP-PCR analysis. (D) Western blot analysis of RBBP4 and RBBP7 in EedKO and Suz12KO ES cells. BAF60 served as a loading control. (E) The binding of distribution of RBBP4 within -5/+5 kb of the center of RBBP4-bound regions in the absence of EED or SUZ12. Heatmaps cover 4140 RBBP4 binding regions identified by MACS2 peak calling. (F) Profile plots depicting the enrichment of RBBP4 within -5/+5 kb of the center of 4020 SUZ12 binding regions and K27ac enriched regions.

We and others have found that loss of EED and SUZ12 destabilizes PRC2-core subunits and most ancillary subunits (Figure 3A) (19). However, RBBP4 and RBBP7 are stable in the absence of EED and SUZ12 (Figure 3D) and are also not required for the stability of other PRC2 core subunits (24). These suggests that RBBP4/7 and other PRC2 components can function as separate units prior to assembly into PRC2 holo-complexes on target loci. To determine if EED and SUZ12 are indispensable for RBBP4 association with chromatin, we performed ChIP-seq analysis to profile the distribution and enrichment of RBBP4 on chromatin in *Suz12*KO and *Eed*KO ESCs. We found that RBBP4 was able to bind to its target genomic loci in knockout cells, but at a lower level compared to the controls (Figure 3E). Next, we separately quantified RBBP4 binding on SUZ12-marked PRC2 loci and HDAC complex loci marked by H3K27ac. Similar to the reduction observed for all RBBP4 target loci, we found slightly reduced RBBP4 binding at SUZ12-, but not H3K27ac-marked loci upon EED depletion, and more striking reduction of RBBP4 binding in the absence of SUZ12 (Figure 3F). Thus, PRC2 holo-complexes help to maintain stable binding of RBBP4 to chromatin. Altogether, these data with the failure of SUZ12 binding to PRC2 target loci due to RBBP4 disruption suggest that RBBP4’s presence at PRC2 loci could serve as a docking site for PRC2 assembly on target chromatin.

### 4. RBBP4 disruption reshapes the genomic landscape of H3K27ac

Since RBBP4 exists in different chromatin modifying complexes, we clustered binding loci into three groups based on the enrichment patterns of H3K27me3, H3K27ac, HDAC1, and p300 (Figure 4A). One cluster (cluster_2) showed strong H3K27me3 signals, indicative of them being PRC2 target loci. This cluster also exhibited enrichment of HDAC1 with concomitant depletion of H3K27ac. However, PRC2 and HDAC1-containing complexes did not interact with each other (Figure S3). We postulated that in these regions, HDAC1 prevented assembly of activating histone acetylation complexes, while H3K27 residues are methylated to maintain a repressive chromatin state. The other two RBBP4 clusters were marked by high and medium levels of H3K27ac and p300 respectively, even though they were also enriched with HDAC1 (Figure 4A). In interphase nuclei, acetylation and deacetylation are dynamic interactions found at specific sites, as well as within large regions in the chromatin to delineate formation and/or organization of euchromatin and heterochromatin domains (30). The colocalization of histone deacetylase HDAC1 and histone acetylase p300 suggests that these genomic regions experience dynamic changes in H3K27ac levels through acetylation and deacetylation processes.

**Figure 4.**
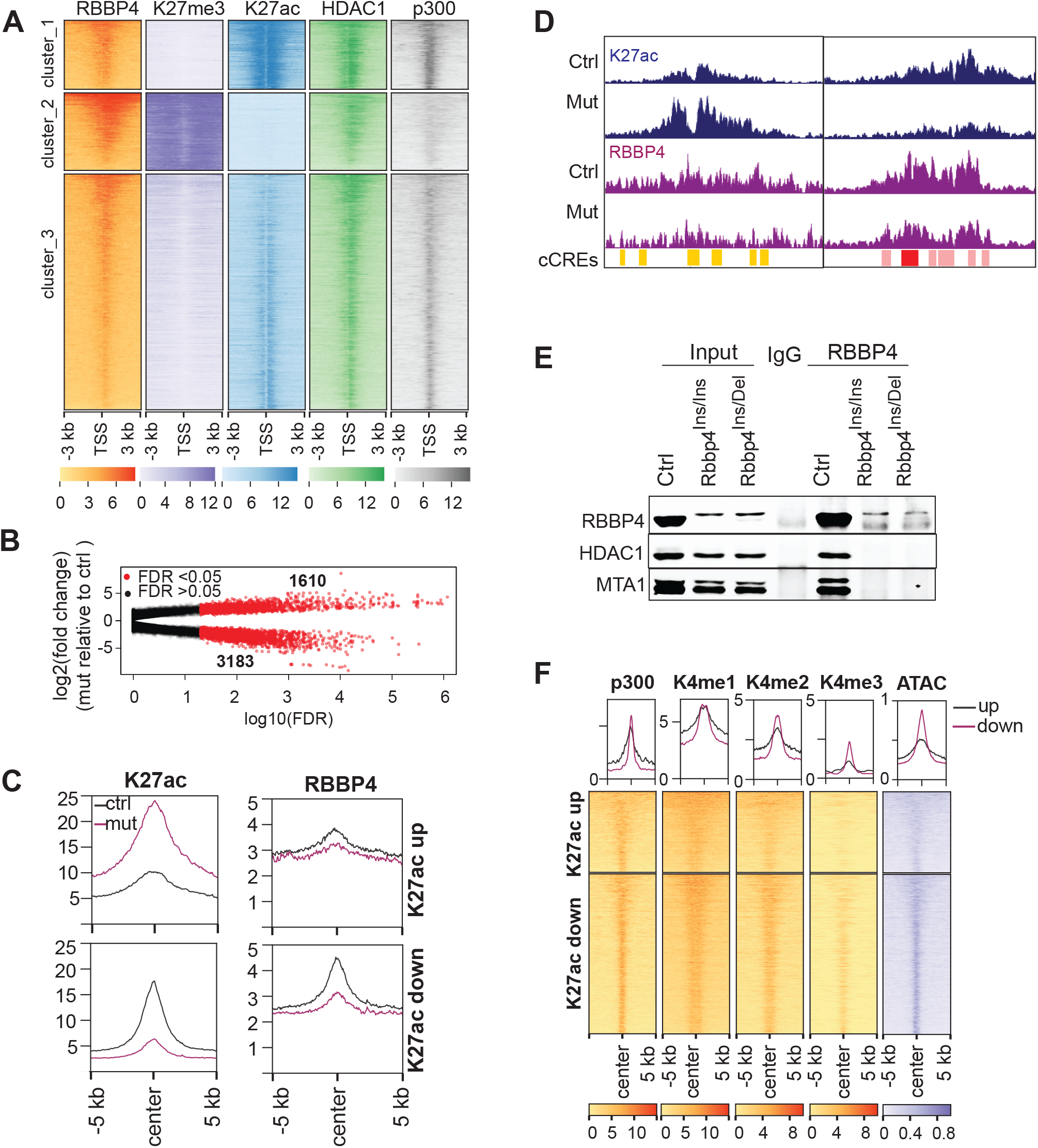
RBBP4 controls acetylation levels of H3K27 on its target chromatin. (A) Clustering RBBP4 binding sites based on methylation and acetylation status of H3K27 within -3/+3 kb of transcription start sites. (B) Comparison of H3K27ac enrichment on its marked genomic regions between the controls and Rbbp4 mutants. Each dot represents a genomic region with H3K27ac binding. (C) The distribution and enrichment of RBBP4, H3K27me3, SUZ12, and HDAC1 within -5/+5 kb of the center of marked by H3K27ac. The genomic loci included for analysis have increased or decreased H3K27ac due to Rbbp4 mutations. (D) Genome browser images of representative genomic loci with increased or decreased H3K27ac incorporation and Rbbp4 binding due to Rbbp4 mutations. The regions are enriched with candidate cis-regulatory elements (cCREs). Yellow or orange bars indicate enhancers, red bars indicate promoters, and pink bars indicate the enrichment of H3K4me3. (E) Co-immunoprecipitation and Western blot analysis for assessing the interaction between endogenous RBBP4 mutants and HDAC1 or MTA1. (F) Heatmaps depicting the genomic regions which had altered H3K27ac in Rbbp4 mutants were marked by enhancer signatures.

Since RBBP4 is the core component of several histone deacetylase complexes, it is expected that disruption of RBBP4 would result in an increase of H3K27ac on chromatin. Differential binding analysis of H3K27ac in E14 and *Rbbp4* mutant ESCs did reveal numerous loci (1610) with increased H3K27ac. However, to our surprise, there were approximately two-fold more (3183) genomic regions with decreased H3K27ac levels (Figure 4B). RBBP4 binding was reduced for both up- and down-regulated H3K27ac loci in the mutants (Figure 4C). Typically, those genomic regions with alternated H3K27ac or RBBP4 bore features of cis-regulatory elements such as enhancers and promoters (Figure 4D).

In probing the ability of the histone deacetylase complex members to interact, we found that the incorporation of mutant RBBP4 into HDAC complexes containing HDAC1 and MTA1 was barely detected (Figure 4E); however the interaction between complex members MTA1 and HDAC1 was maintained in *Rbbp4* mutant cells (Figure S4A). To test whether the reduction of mutant RBBP4 in histone deacetylase complexes was due to its failure to interact with other subunits, we utilized epitope-tagged RBBP4 and HDAC1 for co-immunoprecipitation assays. Similar to wild type RBBP4, mutants were incorporated into HDAC1-containing complexes (Figure S4B). This suggests that RBBP4 plays complex roles in regulatory histone deacetylation dynamics, and its presence in HDAC-containing complexes is important for regulating the activity of the holo-complexes.

We found protein levels of p300, a histone acetylase specialized for H3K27ac, was strikingly decreased in the mutants (Figure S4C), which may explain the extensive reduction of H3K27ac levels in mutant cells. However, p300 mRNA levels were only decreased by about 10% in *Rbbp4* mutants compared to the control (Figure S4D), suggesting possible involvement of RBBP4 in stabilizing p300 protein. However, no physical interaction between p300 and RBBP4/HDAC1/MTA1 was detected (Figure S4C). One future question is based on how the crosstalk between H3K27ac writer and eraser complexes coordinate to regulate acetylation levels.

To characterize the chromatin signature of genomic loci with increased and decreased levels of H3K27ac in the mutants, we retrieved ChIP-Seq data for H3K4me1, H3K4me2, H3K4me3, p300, and ATAC-seq in mouse ESCs from GEO and examined the distribution and enrichment of these features around H3K27ac regions. H3K27ac combined with H3K4 methylation, p300, and open chromatin indicated by ATAC are used to predict cell type-specific enhancers (31). The colocalization of altered H3K27ac loci with these enhancer marks (Figure 4F) suggest that mutations of RBBP4 can impact enhancer activity, leading to gene transcription deregulation.

### 5. RBBP4 plays crucial roles in transcriptional regulation via H3K27 acetylation

RNA-seq analysis performed to assess gene expression changes due to RBBP4 disruption revealed that RBBP4 mutations dramatically altered transcriptional profiles. In *Rbbp4^Ins/Ins^* 3637 genes were significantly (P<0.01) upregulated while 3642 were downregulated relative to control. A similar proportion of genes were mis-regulated in, *Rbbp4^Ins/Del^*, with 4022 upregulated and 3954 downregulated. For those differentially expressed genes which had more than a 2-fold change in transcription and a mean read count of more than 20, there are 762 and 919 upregulated, and 936 and 1066 downregulated genes respectively in *Rbbp4^Ins/Ins^* and *Rbbp4^Ins/Del^* (Figure 5A). These differentially expressed genes showed similar transcriptional patterns between the two mutants (Figure 5B), and included 691 upregulated and 828 downregulated genes (fold change > 2 and read count >20) that were shared by the two mutants (Figure S5A). Such transcriptional mis-regulation is consistent with the abnormal changes in H3K27ac and H3K27me3 that occur in mutant cells.

**Figure 5.**
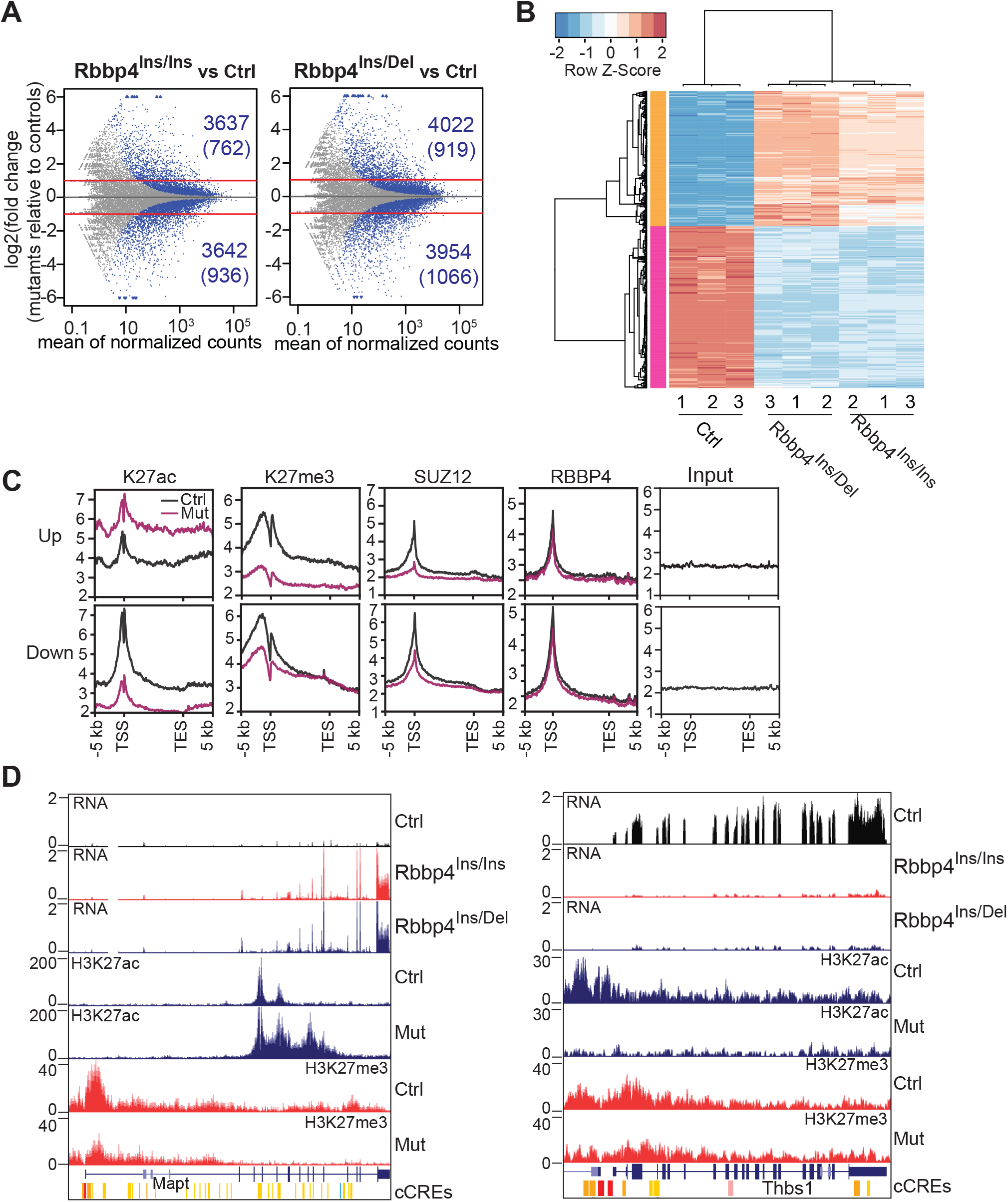
RBBP4 is a master transcriptional regulator through adjusting the levels of H3K27 acetylation. (A) MA-plot shows differential gene expression due to RBBP4 dysfunction. The blue dots represent genes with significant changes in transcription (p< 0.01). Red lines indicate two-fold changes. (B) A heatmap of the top 1000 differentially expressed genes. (C) Comparison of the enrichment of H3K27ac, H3K27me3, RBBP4, and SUZ12 over gene bodies of transcriptionally mis-regulated genes between the controls and Rbbp4 mutants. (D) Genome browser images of representative genes whose activation and silencing is coordinated with gain and loss of H3K27ac on cis-regulatory elements.

In contrast to moderate gene expression changes in *Eed*, *Ezh2*, or *Suz12*-depleted mouse ESCs, RBBP4 disruption led to transcription changes at a much larger scale (Figure S5A,B) (24, 32, 33). In *Eed* knockout ESCs, there were 580 and 368 genes whose transcription increased and decreased by more than 2-fold respectively, compared to the 967 and 1207 genes with observed changes in RBBP4 mutants. There were only 97 upregulated and 83 downregulated genes shared by *Eed* knockout and *Rbbp4* mutants. Unexpectedly, 115 deregulated genes in the *Eed* knockout were inversely regulated with dramatic changes in *Rbbp4* mutants (Table S1). For example, the expression of *Frem2* and *Gsta3* are up- and down-regulated, respectively, in *Eed* knockout ESCs but were oppositely regulated in *Rbbp4* mutants (Figure S5C). We postulate that except for PRC2-mediated transcriptional repression, RBBP4 controls gene activity through histone deacetylation complexes.

We examined the enrichment of H3K27ac, H3K27me3, RBBP4, and SUZ12 on commonly de-regulated genes in the two *Rbbp4* mutants (Figure 5C). The incorporation of H3K27ac was markedly increased across the up-regulated gene bodies and decreased for the downregulated genes in the mutants. H3K27me3 and SUZ12 levels were lower on both up-and down-regulated genes because disruption of RBBP4 caused broad reduction of H3K27me3 and SUZ12 binding. For instance, *Mapt* expression increased by 1.4-fold upon depletion of H3K27me3 in *Eed* KO ES cells (32), while in contrast, *Mapt* expression increased by 14.4-fold upon gaining H3K27ac that spans predicted cis-regulatory elements, concomitant with only a partial loss of H3K27me3 in *Rbbp4* mutant ESCs (Figure 5D). Thus, enhanced activation of *Mapt* in *Rbbp4* mutants was primarily related to the acquisition of H3K27ac rather than a loss of H3K27me3. We also observed that *Thbs1* was silenced with loss of H3K27ac upstream of its promoter, but not via gaining repressive H3K27me3 (Figure 5D). Together, these data shows that RBBP4-mediated metabolism of H3K27ac on chromatin plays a key role for controlling gene activity.

### 6. RBBP4 regulates gene activity by controlling H3K27 acetylation on enhancers and super-enhancers

To explore the mechanisms by which RBBP4 disruption alters H3K27ac to perturb gene expression, we characterized the genomic features of those H3K27ac marked regions. Using the ENCODE mouse cis-regulatory elements database as a reference, we found that both up- and down-regulated H3K27ac loci are significantly enriched with enhancer-like signatures (5836 and 7165 respectively) (Table 1). Selected loci also cover many promoters and other cis-regulatory elements as defined by DNase hypersensitivity, H3K4me3 and CTCF binding (Table 1). We found RBBP4 associated with sites normally devoid of H3K27me3 and SUZ12 but marked with H3K27ac thereby resembling putative enhancers (Figure 6A). These sites displayed both increased and decreased H3K27ac concomitant with reduced RBBP4 binding in mutant ESCs, suggesting that RBBP4 is involved in regulating enhancer activity.

**Figure 6.**
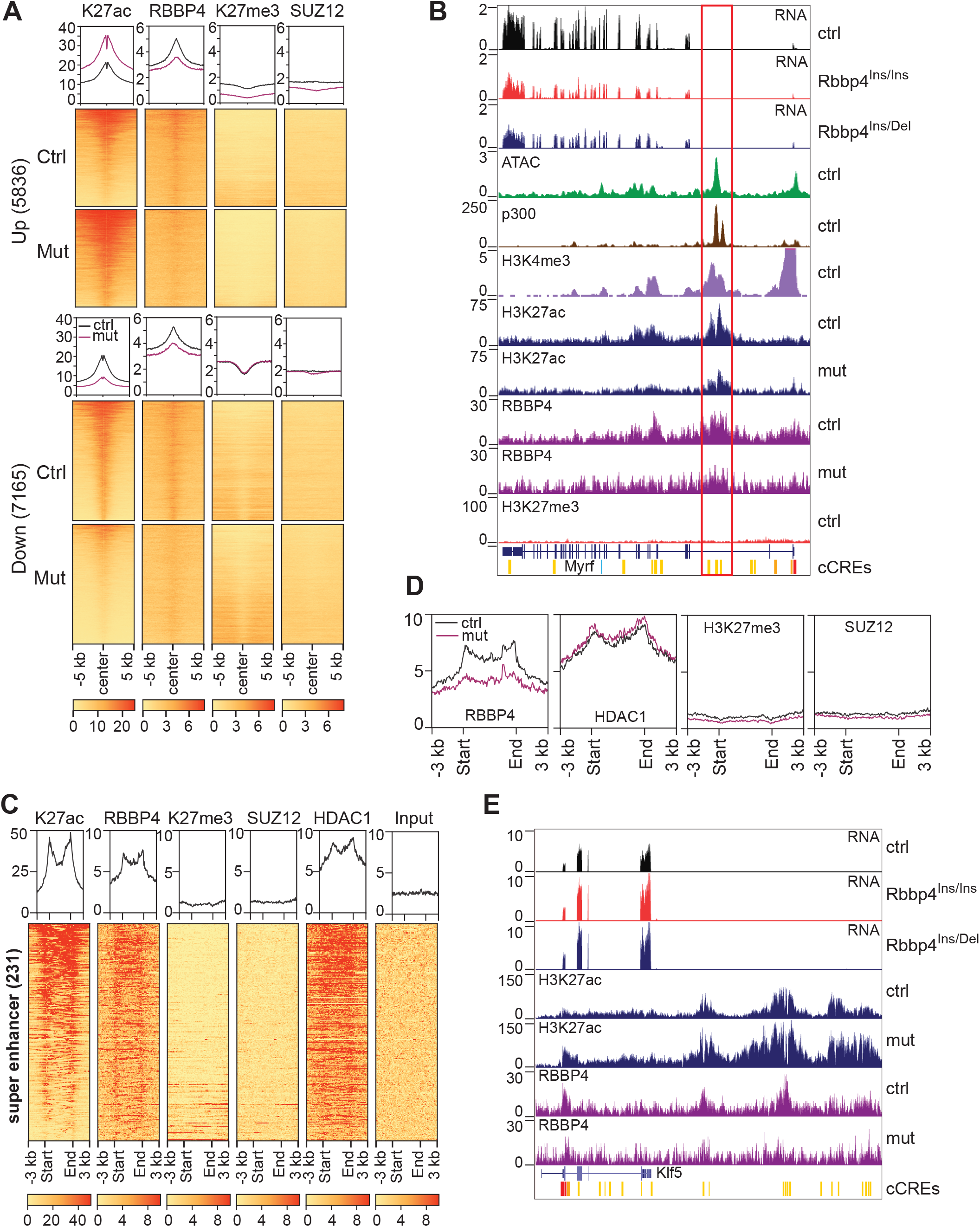
RBBP4 regulates gene activity by controlling H3K27 acetylation on enhancers and super-enhancers. (A) H3K27ac, RBBP4, H3K27me3 and SUZ12 enrichment at putative enhancers. Heatmaps are centered within -5/+5 kb of K27ac enriched loci. (B) Genome browser image of Myrf locus. Red rectangle indicates Myrf enhancer region. (C) RBBP4 and HDAC1 enrichment on super-enhancers. The heatmaps cover the regions within 3 kb upstream and downstream of super-enhancers. (D) Plot profile depicting RBBP4 enrichment on super-enhancers in control (black line) and Rbbp4 mutant (magenta line) ESC’s. (E) Genome browser images showing increased H3K27ac on a putative Klf5 enhancer in Rbbp4 mutant was related to upregulation of this gene.

**Table 1.** The genomic regions with altered H3K27ac in the mutants are enriched with predicated cis-regulatory elements

| **Genomic feature** | **Up** | **Down** |
| --- | --- | --- |
| **K27ac regions** | 1610 | 3183 |
| **promoter** | 494 | 1074 |
| **proximal enhancer** | 1621 | 2694 |
| **distal enhancer** | 4215 | 4471 |
| **super enhancer** | 59 | 17 |
| **regions with other CRE** | 1053 | 2286 |

We examined the impact of H3K27ac and RBBP4 levels on the activity of validated enhancers. In mouse oligodendrocytes, one of *Myrf*’s enhancers was found to be in an intron (34). In mouse ESCs, that enhancer is also in an active state as indicated by several features including H3K27ac, H3K4me3 and p300 chromatin marks, an open chromatin ATAC signature, and high *Myrf* transcription levels (Figure 6B). In *Rbbp4* mutant cells, RBBP4 binding and H3K27ac accumulation were decreased around the *Myrf* enhancer, resulting in transcriptional suppression (Figure 6B). Thus, RBBP4 can control gene activity through regulating H3K27ac levels on enhancers.

Super-enhancers are marked by clusters of H3K27ac and function to maintain high expression of cell-type specific genes for cell identity maintenance (35). We found that dysregulated H3K27ac was enriched on super-enhancers in E14 ESCs. Among the 231 super-enhancers present in mouse ESCs (36), 76 super-enhancers underwent H3K27ac changes in RBBP4 mutants (Table 1). RBBP4 and HDAC1 bind to super-enhancer regions and exhibit similar distribution patterns as H3K27ac (Figure 6C). Compared to E14 cells, less RBBP4 was bound to super-enhancers in RBBP4 mutants, but this did not affect the binding of HDAC1 at these regions (Figure 6D).

We next explored individual contexts of relevant gene modulation in ESCs as RBBP4 has been shown to be involved in the maintenance of mouse ESC pluripotency (24). We focused on *Klf5* since it is a regulator of ESC pluripotency (37) and prevents differentiation toward mesoderm (38). Given that *Klf5* is important for controlling and defining ESC identity, we predicted that the Klf5 locus and its upstream region, which are broadly marked by H3K27ac (Figure 6E), is a super-enhancer. Here, we observed H3K27ac levels across this region were elevated along with decreased binding of RBBP4 in mutant cell lines (Figure 6E). Correspondingly, the transcription of *Klf5* was enhanced (Figure 6E). Given these specific examples and genome-wide data, our analysis indicates that RBBP4 adjusts H3K27ac levels at super-enhancers to finely tune gene expression necessary for stem cell homeostasis and differentiation.

## Discussion

In mouse ESCs, H3K27me3 is highly enriched in regulatory regions of key developmental genes and helps to maintain these genes in a silenced state (39). These histone marks are transmitted to daughter cells during cell proliferation for cell identity maintenance. PRC2 localizes to genomic loci marked by H3K27me3 but not H3K27me1 and H3K27me2, suggesting that PRC2 forms stable complexes on chromatin to catalyze H3K27me3 modification (1). It has been reported that CpG-rich DNA elements, repressive transcriptional states, chromatin-binding proteins, DNA modifications, histone modifications, and noncoding RNA mediate the interaction between PRC2 and chromatin (1, 40). However, these have not fully explained how PRC2 subunits are recruited to site-specific chromatin regions for H3K27me3.

As a core subunit of PRC2, RBBP4 is a histone chaperone protein that directly binds to histones H3 and H4 (20). It is a core component of multiple chromatin-modifying complexes and is essential for viability, which makes it difficult to study specific roles in PRC2-related histone methylation versus HDAC-related acetylation for example. These data led us to investigate whether RBBP4 could mark PRC2 target chromatin for propagation of H3K27me3. Using the CRISPR-Cas9 system to introduce mutations in *Rbbp4*, we generated viable ESC lines with expression of altered RBBP4 proteins. These mutations resulted in striking changes in the genomic enrichment and distribution of both H3K27me3 and H3K27ac, as well as a reshaped transcriptional profile. First, we found that RBBP4 can guide the assembly of PRC2 subunits on target chromatin to for trimethylation of H3K27. Additionally, RBBP4 participates in both acetylation and deacetylation of H3K27, where it controls gene activity by regulating acetylation levels on cis-regulatory elements, especially enhancers.

Regarding PRC2, we found that RBBP4 protein stability is maintained independent of other core subunits, along with its ability to bind PRC2 target chromatin in the absence of EED and SUZ12. We also found that loss of RBBP4 did not affect other PRC2 subunits, confirming that these complex members can independently exist in separate complexes prior to assembly on target chromatin. However, disruption of RBBP4 impairs the recruitment of SUZ12 and EZH2 to PRC2 target loci, leading to a decrease in H3K27me3. Therefore, pre-existence of RBBP4 on nucleosomes could serve as a cue for recruiting other PRC2 subunits to restore H3K27me3 on newly synthesized histones.

DNA replication leads to disassembly of nucleosomes into H3-H4 tetramers and H2A-H2B dimers (41, 42), with most parental histone (H3-H4)2 tetramers being maintained in close vicinity to their original locus (43, 44). To reproduce a similar chromatin environment on new DNA, histones and perhaps other chromatin-bound factors are transferred from parental strands to daughter strands (45). It is possible that RBBP4 binds to H3-H4 tetramers during nucleosome disassembly to direct other subunits of PRC2 towards new histones in their vicinity to maintain faithful transmission of epigenetic information.

A recent study demonstrated that SUZ12 bond to PRC2 target regions independently of EED and EZH1/2 where it is essential for guiding PRC2 to its correct genomic loci (18). We found that RBBP4 disruption interfered with SUZ12 recruitment to PRC2 loci, suggesting that RBBP4 association with chromatin is also an early event in PRC2’s assembly on target genomic regions. SUZ12-RBBP4 complex guides incorporation of ancillary subunits to form different PRC2 subcomplexes on chromatin (23, 46), and we observed that loss of SUZ12 caused decreased accumulation of RBBP4 on PRC2 target chromatin. We therefore propose that RBBP4 and SUZ12 coordinate to initiate PRC2 assembly on specific genomic sites.

Compared to a developmental arrest at the gastrulation stage due to depletion of EED (7), SUZ12 (47), and EZH2 (8), RBBP4 ablation resulted in more severe phenotypes such as pre-implantation lethality and failure of inner cell mass outgrowth (ICM) (48). This indicates that non-PRC2 related functions such as RBBP4-mediated histone deacetylation processes are crucial for ICM proliferation and lineage commitment. RBBP4 coexists with HDAC1 in deacetylase complexes that broadly deacetylates lysines (49). One of these, the NuRD complex, where RBBP4 along with MTA1 and HDAC1 are core subunits, was found to bind on a genome-wide scale to nearly all active enhancers and promoters in ESCs (50). Based on these results, and the knowledge that H3K27ac is enriched in cis-regulatory elements to finely regulate gene activity, in this study we also focused on the impact of RBBP4 on acetylation status. As expected, RBBP4 disruption caused extensive elevation of H3K27ac on enhancers, which is further indicative of the importance of RBBP4-containing complexes in gene activation states.

Mutant RBBP4 was insufficiently incorporated into complexes containing HDAC1 and MTA1, a possible mechanism behind our observations of impaired deacetylation activity on enhancers and promoters. The NuRD complex contains four RBBP4 subunits, which could provide a structural basis for optimization of HDAC1/2 catalytic activity. NuRD has a general affinity for open chromatin regions associated with transcriptional activity and fine-tunes gene expression rather than acting as categorical “on–off” switches (51). As *Rbbp4* mutations did not arrest ESC proliferation, it is possible that the mutations did not disrupt the functions of other RBBP4-containing complexes, such as Sin3, which are essential for ESC self-renewal (52). However, it is known that under self-renewal conditions, NuRD binds to a subset of pluripotency genes (Klf4, Klf5, and Tbx3) where it confines their expression (53). Consistent with this, RBBP4 mutant ESCs had numerous upregulated genes, including pluripotency genes, which are surrounded by increased levels of H3K27ac. Altogether, this suggests that RBBP4 helps to maintain pluripotency regulatory networks through regulating NuRD’s deacetylation activity on H3K27.

Regarding crosstalk between histone modifications, it has been established that RBBP4 is involved in the regulation of acetylation and methylation on the same lysine residues of histone H3 for transcriptional repression. The genomic distribution of these two modifications is mutually exclusive and we did not observe interactions between PRC2 and NuRD, indicating that methylation and deacetylation of H3K27 are independent events in gene repression. In support of this, we found only a few overlapping dysregulated genes between *Rbbp4* disruption and *Eed* knockout in ESCs. Compared to ablation of *Eed*, *Suz12*, and *Ezh2*, RBBP4 disruption caused more widespread transcriptional deregulation (24, 32, 33). We also found a subset of genes that were discordantly regulated between *Eed* and *Rbbp4* mutants with respect to loss or gain of H3K27ac. Together, these data suggest that RBBP4 controls gene activity primarily through regulating H3K27ac levels.

A previous study showed sparse increases of H3K27ac along the genome as a consequence of PRC2 loss, and suggested chromatin hyperacetylation rather than specific loss of repressive control at target genes leads to the early developmental failure induced by PRC2 disruption (54). In mouse ESCs, *Hox* cluster genes are highly enriched with H3K27me3, but nearly all these genes did not experience transcriptional activation due to loss of H3K27me3. H3K27me3 marks poised chromatin and helps to maintain a transcriptionally silenced state, which is critical for spatiotemporal activation of cell-type specific genes during development. In contrast, H3K27ac is a more potent gene expression activator that actively and dynamically regulates transcriptional levels.

This leads us to propose a model for RBBP4’s function in regulating methylation and deacetylation/acetylation of H3K27. On one hand, RBBP4 consistently binds to parental histones during DNA replication and guides the assembly of other PRC2 subunits to adjacent newly synthesized histones for maintaining the genomic landscape of H3K27me3 during cell proliferation (Figure 7A). One the other hand, RBBP4 facilitates the deacetylase activity of HDAC complexes for efficient removal of acetyl group on H3K27. Meanwhile, RBBP4 also promotes H3K27ac through maintaining p300 levels. Altogether therefore it is possible that RBBP4 assists in controlling H3K27ac levels on cis-regulatory elements for exquisite programming of the transcriptome with regards to cellular context. Better understanding of the molecular mechanisms of RBBP4 will help to elucidate its functional specificities in chromatin regulation and gene expression. In addition to our findings in this study that are relevant to developmental biology, pharmacological interventions to manipulate H3K27me3 and H3K27ac are increasingly explored for cancer therapy. As RBBP4 sits at the intersection of the H3K27 epigenome, it warrants this interest, as it may hold broad implications for chemotherapy applications.

**Figure 7.**
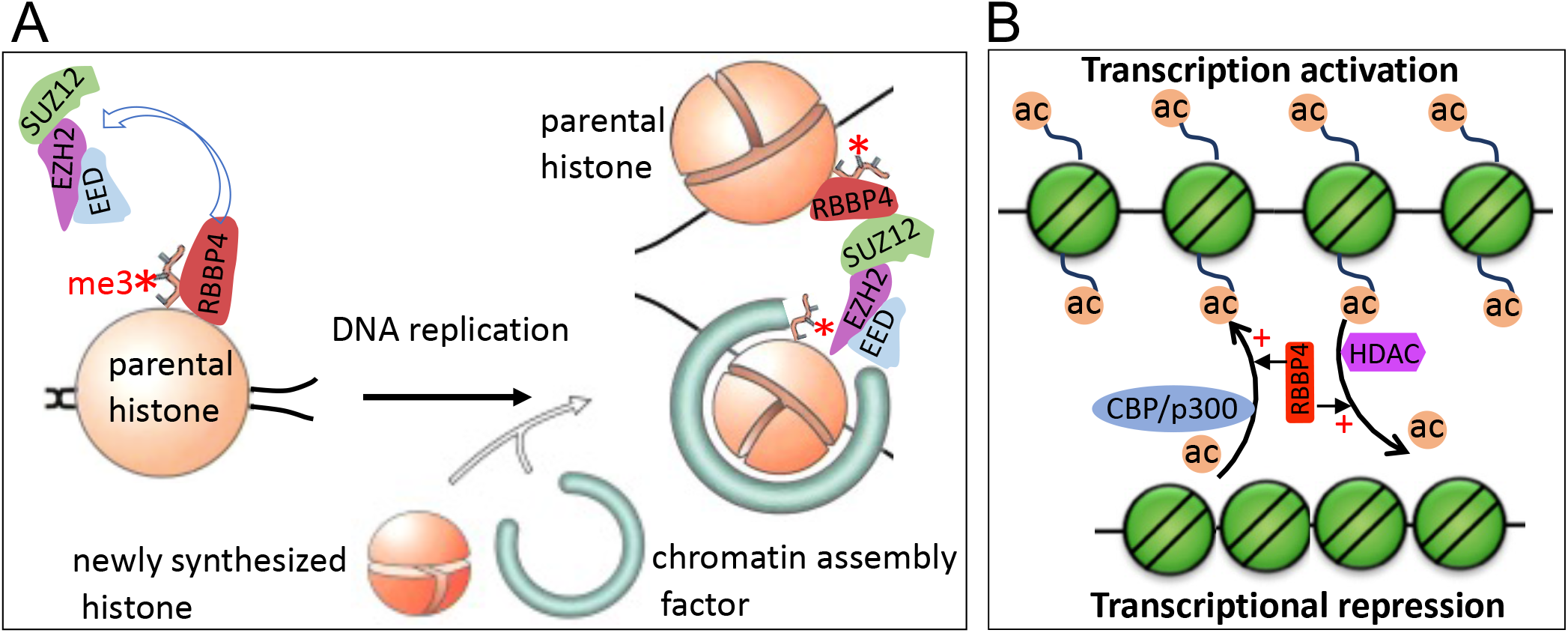
A model for RBBP4’s function in regulating methylation and deacetylation/acetylation of H3K27. (A) RBBP4 consistently binds to parental histones during DNA replication and guides the assembly of other PRC2 subunits to adjacent newly synthesized histones for trimethylation of H3K27. As a result, genomic landscape of H3K27me3 is maintained during cell proliferation. (B) The presence of RBBP4 in HDAC complexes can facilitate deacetylase activity for efficient removal of acetyl groups on H3K27. Meanwhile, RBBP4 can promote H3K27ac through maintaining p300 levels. Thus, RBBP4 controls H3K27ac levels on cis-regulatory elements to finely tune transcription levels.

## Materials and methods

### Sample-size estimation

The experiments in this study were performed with cell lines, and, because there was no biological variation to account for, no power analysis was performed in advance. Also, since all experiments were done using cell-lines, all replication was technical in nature. That said, Western blots used 1 or more technical replicates per cell line. These provided qualitative results, and when multiple technical replicates were used, they served only for confirmation. ChIP-seq used one cell line per antibody. Because the results were verified with ChIP-qPCR, ChIP-seq was performed without additional technical replicates. RNA-seq was performed in triplicate to provide power for detecting differential gene expression. We decided on three technical replicates for RNA-seq based on previous experience with RNA-seq and its ubiquity in the field. There were no data outliers and no experiments were excluded.

### Genome editing of Rbbp4 and Rbbp7 by CRISPR-Cas9

sgRNAs targeting *Rbbp4* and *Rbbp7* were cloned into eSpCas9(1.1) (Addgene, Cat. No. 71814) using a Golden Gate assembly cloning strategy (55). The modification of *Rbbp4/7* genes in ESCs followed the procedure as described (56). Briefly, 5 x 104 E14 ESCs were cultured on 60 mm dishes for 1 day and then transfected with plasmids expressing Cas9 and sgRNAs, along with a plasmid expressing PGK-PuroR (Addgene, Cat. No. 31937) using the FuGENE HD reagent (Promega) according to the manufacturer’s instructions. The cells were treated with 2 μg/ml puromycin for 2 days and recovered in normal culture medium until ESC colonies grew. Targeted colonies were genotyped by PCR and verified by DNA sequencing and Western blot analysis.

### ChIP-seq analysis

ChIP-seq was performed as described (57) with minor modifications. Five million cells were used per IP. The nuclear membrane was broken by mild sonication instead of passing through 20G needles. ChIP-seq libraries were prepared using the KAPA HyperPrep kit according to manufacturer’s instructions and sequenced on Illumina’s NovaSeq 6000 system. Sequence reads were aligned to genomic sequence (mm10) with Bowtie2 (58). MACSv2 identified ChIP-seq enrichment using the broad peaks model (59). Differential binding analysis were performed using CSAW and significant differences in counts were called at a false discovery rate (FDR)≤0.05 (60). The ChIP-seq experiment for each antibody and cell line was performed once.

### RNA-seq analysis

RNA-seq analysis was performed in triplicate for control and each mutant cell lines. Cells were lysed with the TRIzol reagent (Invitrogen) and total RNA was isolated using the Direct-zol RNA kit (Zymo). Sequencing libraries were prepared using a Kapa mRNA HyperPrep kit as per the manufacturer’s instructions and then sequenced on an Illumina NovaSeq 6000 system (paired end x 50 bp). Sequence reads were aligned to mm10 genomic sequence with STAR (v2.7.3) (61). Aligned reads were counted by HTSeq (62) and differentially expressed genes between controls and mutants were analyzed using DESeq2 (v1.22.2) and significant differences in counts were called at FDR adjusted p-values ≤0.01 (63). A heatmap of top one thousand differentially expressed genes was drawn with Heatmap.2. The RNA-seq experiment was performed once.

### Cell culture

Mouse E14 ESCs were cultured in Glasgow Minimum Essential Medium supplemented with 15% fetal bovine serum, 1.0 mM L-glutamine, 0.1 mM minimal essential medium-nonessential amino acids, 0.1 mM β-mercaptoethanol, and leukemia inhibitory factor. HEK293 were cultured in Dulbecco’s modified Eagle’s medium supplemented with 10% fetal bovine serum, 1.0 mM L-glutamine, 0.1 mM minimal essential medium-nonessential amino acids, 1.0 mM sodium pyruvate, and 0.1 mM β-mercaptoethanol. The cells were transfected with FLAG- or HA-tag expression vectors using calcium phosphate and harvested 48 hours post-transfection for protein extraction. FLAG- or HA-tagged coding fragments of RBBP4, SUZ12, EED, HDAC1, and MTA1 were amplified from their cDNAs and cloned into an expression vector bearing the CAG promoter.

### Protein extraction, histone extraction, immunoprecipitation, and Western blotting

Protein or histones were prepared from ESCs or HEK293T cells for Western blot analysis as described (64). For immunoprecipitation assays, whole cell lysate was prepared in F lysis buffer (20 mM Tris pH 7.9, 500 mM NaCl, 4 mM MgCl2, 0.4 mM EDTA, 2 mM DTT, 20% glycerol, 0.1% NP40, and proteinase inhibitor) and adjusted to 300 mM NaCl by adding dilution buffer (20 mM Tris pH 7.9, 10% glycerol) (65). 200 μg of protein extract was incubated with antibodies for immunoprecipitation. For cell fractionation, E14 ESCs were lysed in CSK buffer (10 mM PIPES pH 6.8, 100 mM NaCl, 300 mM sucrose, 3 mM MgCl2, 1 mM, 0.5% Triton X-100, 1 mM DTT and protease inhibitors) for collection of non-chromatin bound protein, and the remaining pellet incubated in F lysis buffer for preparation of chromatin-bound protein. For Western blot, some experiments were repeated for a high-quality image, but the results were consistent among duplicated experiments.

### Immunofluorescence staining

Immunostaining on mouse ESCs is as described (66). *Eed* KO and *Suz12* KO ES cells were transfected with FLAG-tagged EED or SUZ12 using Xfect™ mESC Transfection Reagent (631320, Clontech) according to manufacturer’s instruction.

### GEO accession

H3K4me1 (GSM1908888), H3K4me2 (GSM881353), H3K4me3 (GSM723017), p300 (GSM2360934), and ATAC-seq (GSE120393).

### Data accessibility

GSE183291, GSE183292, and GSE183298.

## Acknowledgement

We thank all members of the Magnuson Lab for helpful comments on manuscript preparation, and Karl B Shpargel, Prabuddha Chakraborty, Jesse Raab, and Josh Starmer for sharing their scripts for NGS data analysis. This work was supported by National Institutes of Health grants RO1 GM101974 and U42 OD010924.

## Competing Interest Statement

The authors declare no competing interests

---

**Figure S1.**
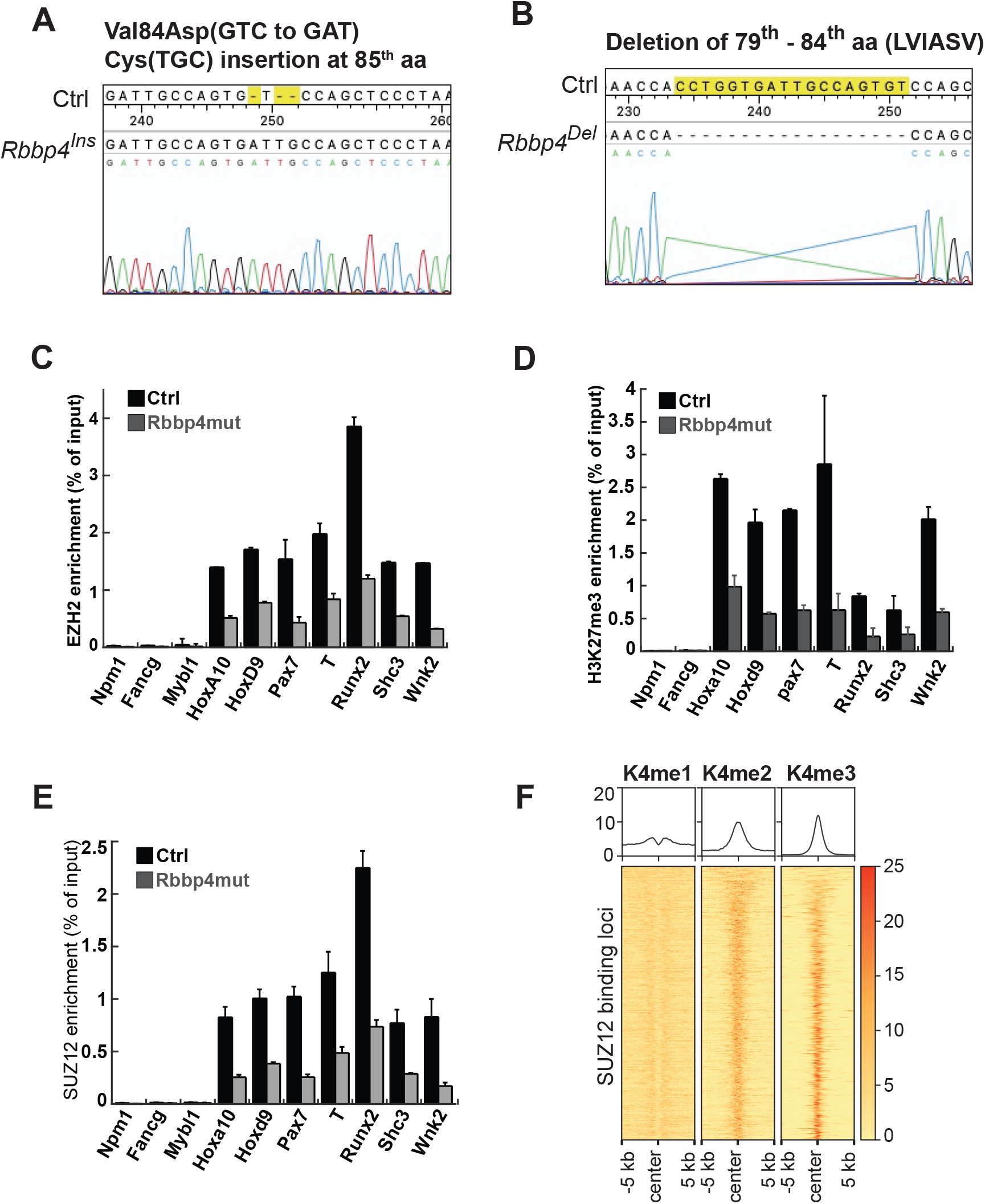
Characterization of Rbbp4 mutant ESC lines by CRISPR-Cas9. (A) Identification of mutations by DNA sequencing in Rbbp4Ins/Ins. (B) Identification of mutations by DNA sequencing in Rbbp4 Ins/Del. (C) ChIP-PCR analysis of EZH2 enrichment on PRC2 target genes. (D) ChIP-PCR analysis of H3K27me3 enrichment on PRC2 target genes. (E) ChIP-PCR analysis of SUZ12 enrichment on PRC2 target genes. (F) Heat maps depicting the enrichment of H3K4 methylation on PRC2 marked loci

**Figure S2.**
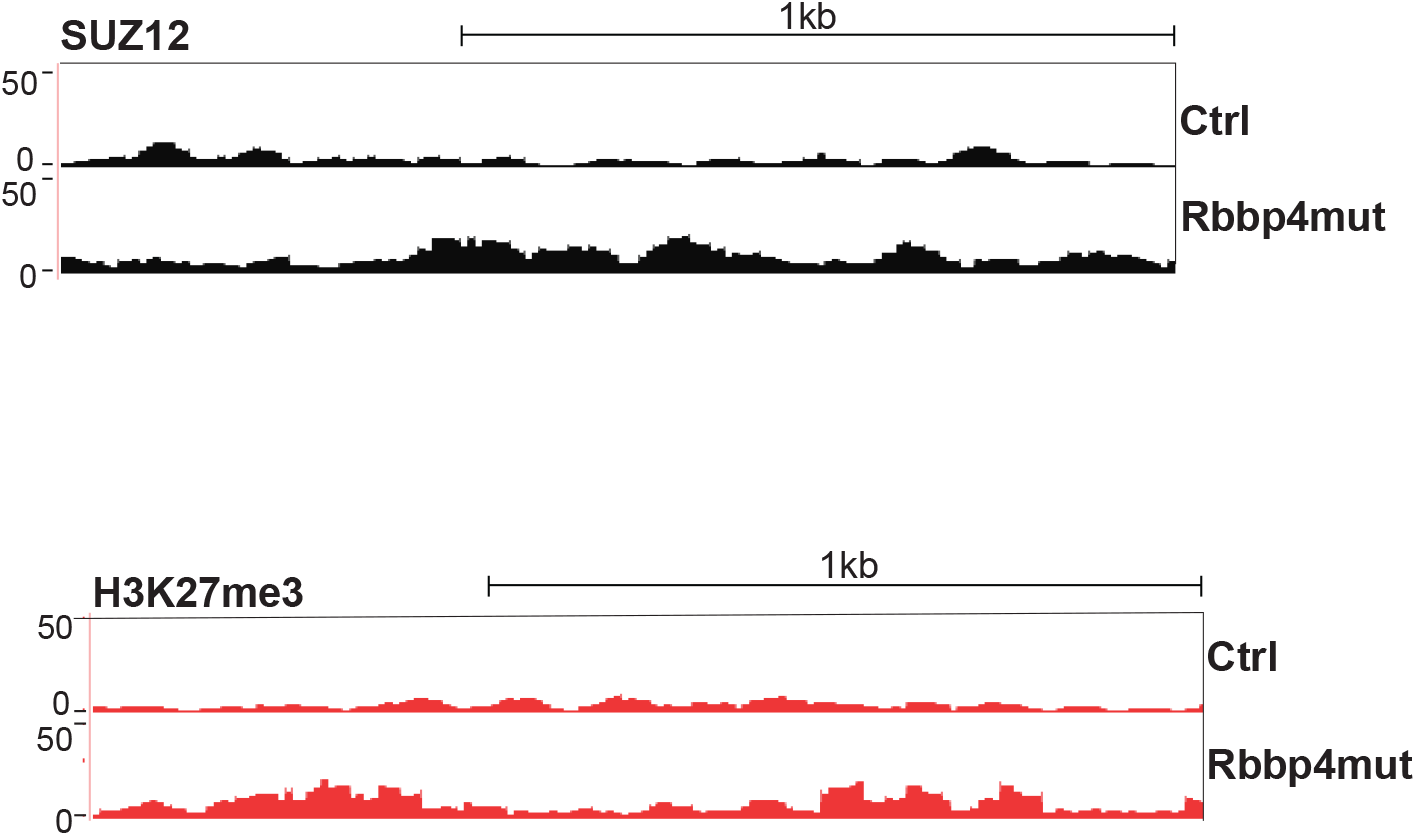
Genome browser image of genomic region with increased SUZ12 and H3K27me3 in Rbbp4 mutant ESCs identified by differential binding analysis.

**Figure S3.**
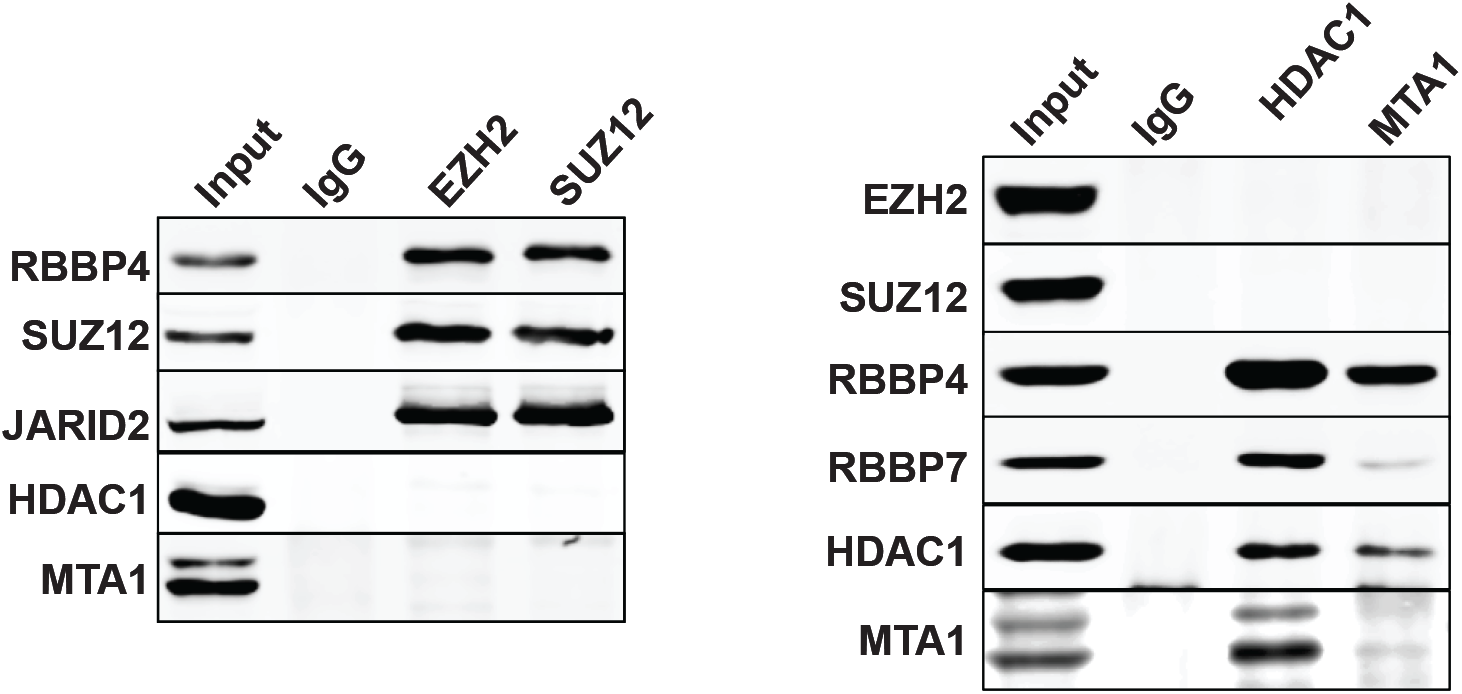
Co-immunoprecipitation and Western blot analysis on the interaction between PRC2 and HDAC1-containing complex.

**Figure S4.**
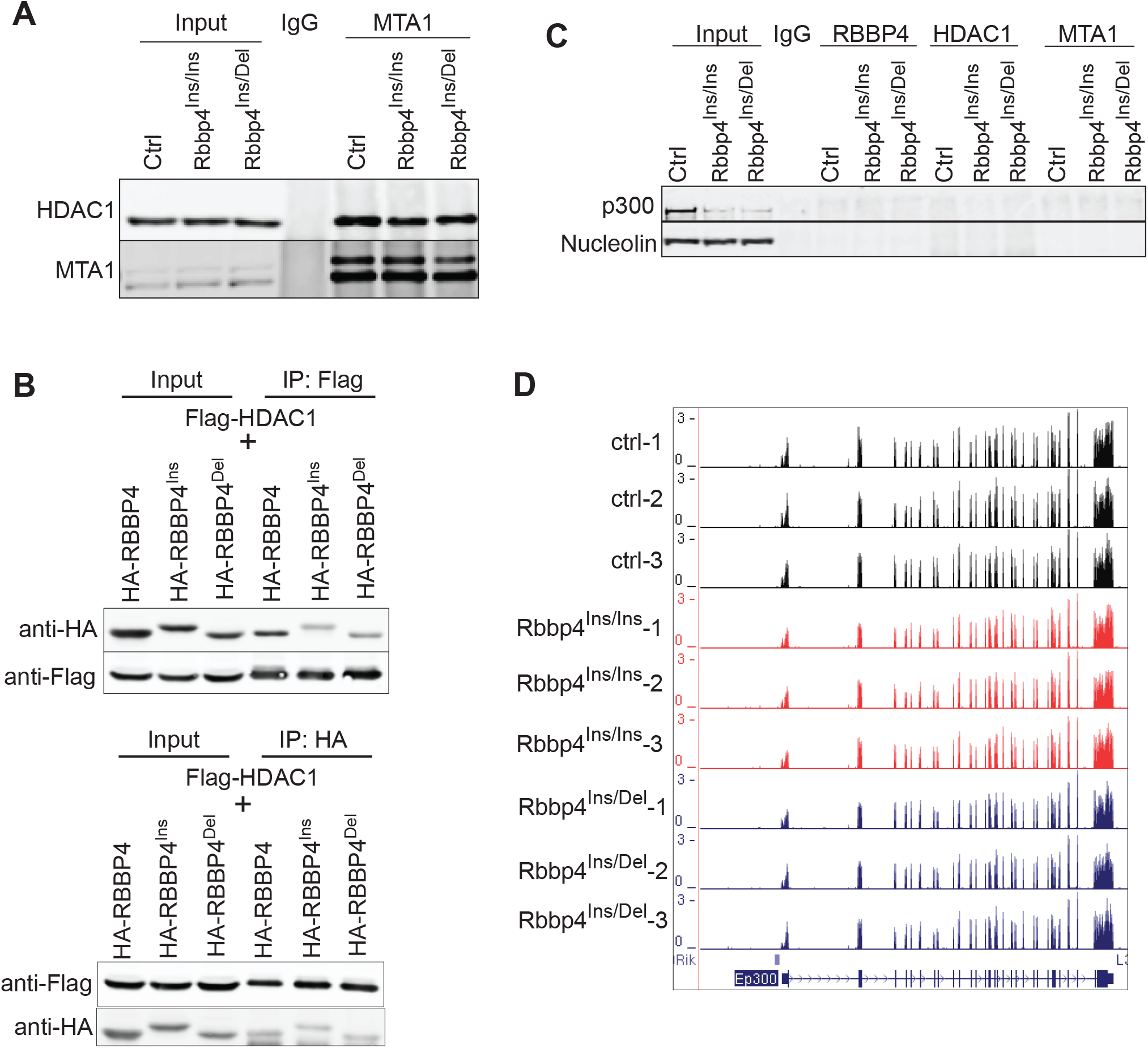
(A) Co-immunoprecipitation and Western blot analysis show that the interaction between HDAC1 and MTA1 was not affected due to RBBP4 disruption. (B) Co-immunoprecipitation and Western blot analysis for verifying the interaction between RBBP4 mutants and HDAC1. HA-tagged and Flag-tagged proteins were expressed in HEK293T cells. (C) Co-immunoprecipitation and Western blot analysis on the interaction between p300 and NuRD complex subunits. (D) UCSC genome browser snapshot of p300 transcriptional levels by RNA-Seq analysis.

**Figure S5.**
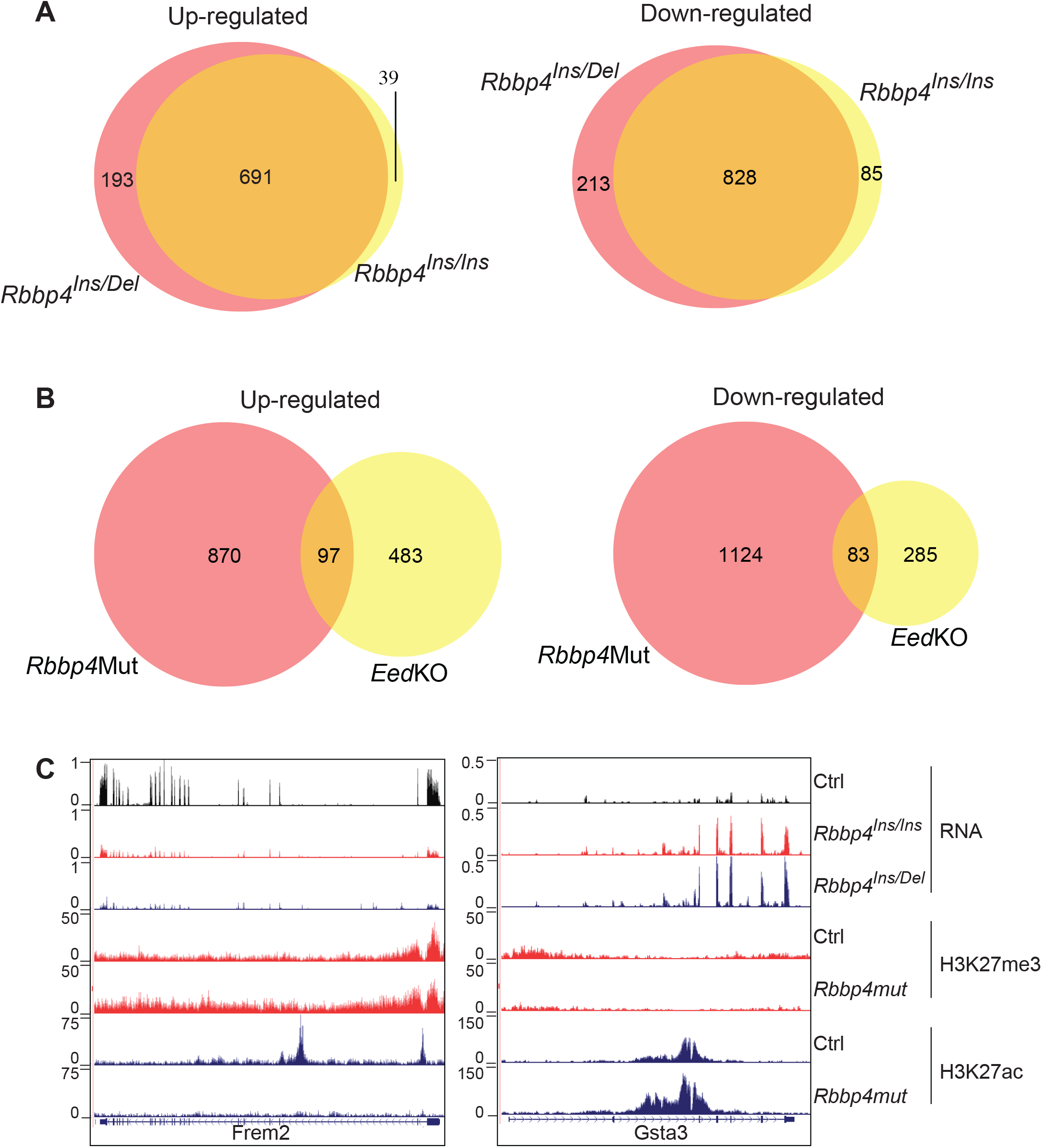
(A) Venn diagrams showing the overlap of differentially expressed genes with more than 2-fold changes between Rbbp4Ins/Ins and Rbbp4Ins/Del ESCs. (B) Venn diagrams showing the overlap of differentially expressed genes with more than 2-fold changes between Rbbp4 mutant and Eed knockout ESCs. (C) Genome browser images of representative genes the transcription of which was reversely deregulated in Rbbp4 mutants and Eed knockout.

**Table S1.**
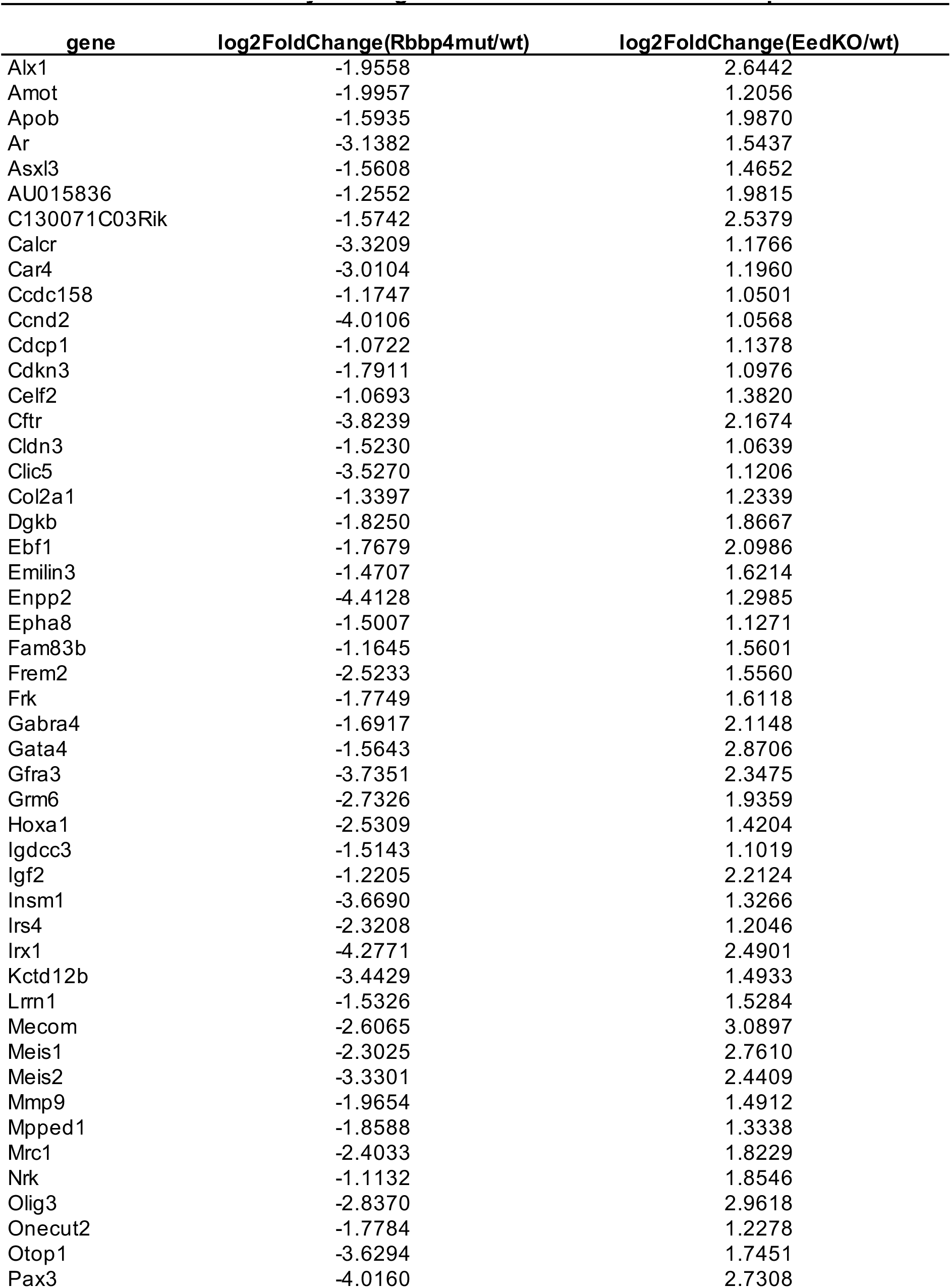
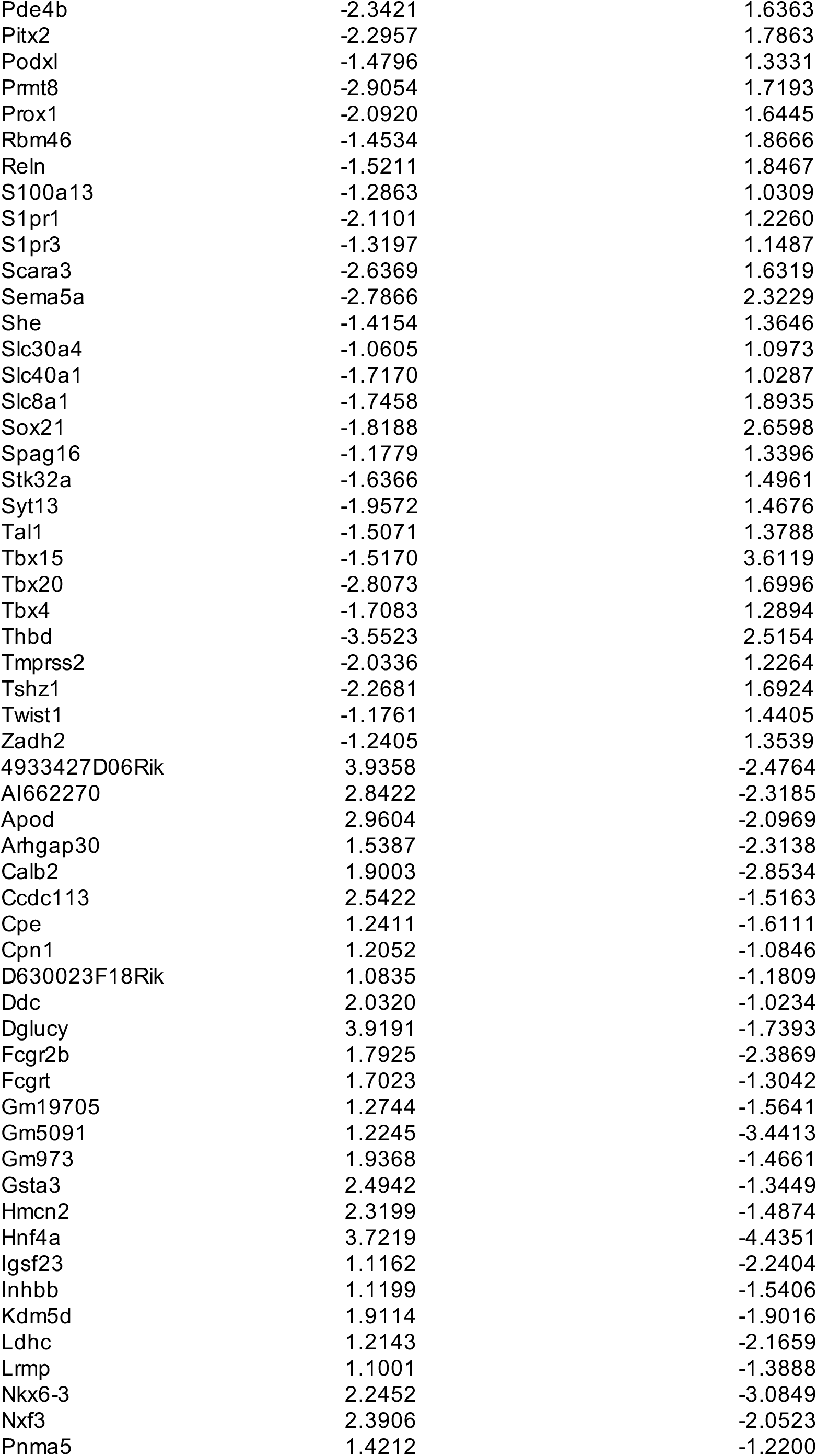
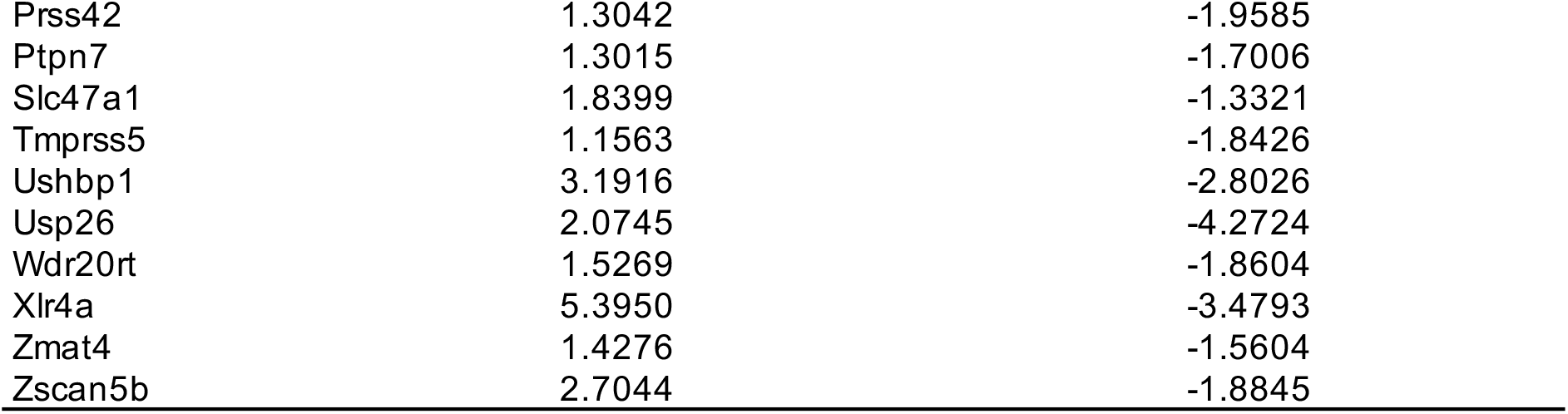
Genes are inversly mis-regulated in Eed knockout and Rbbp4 mutant ESCs

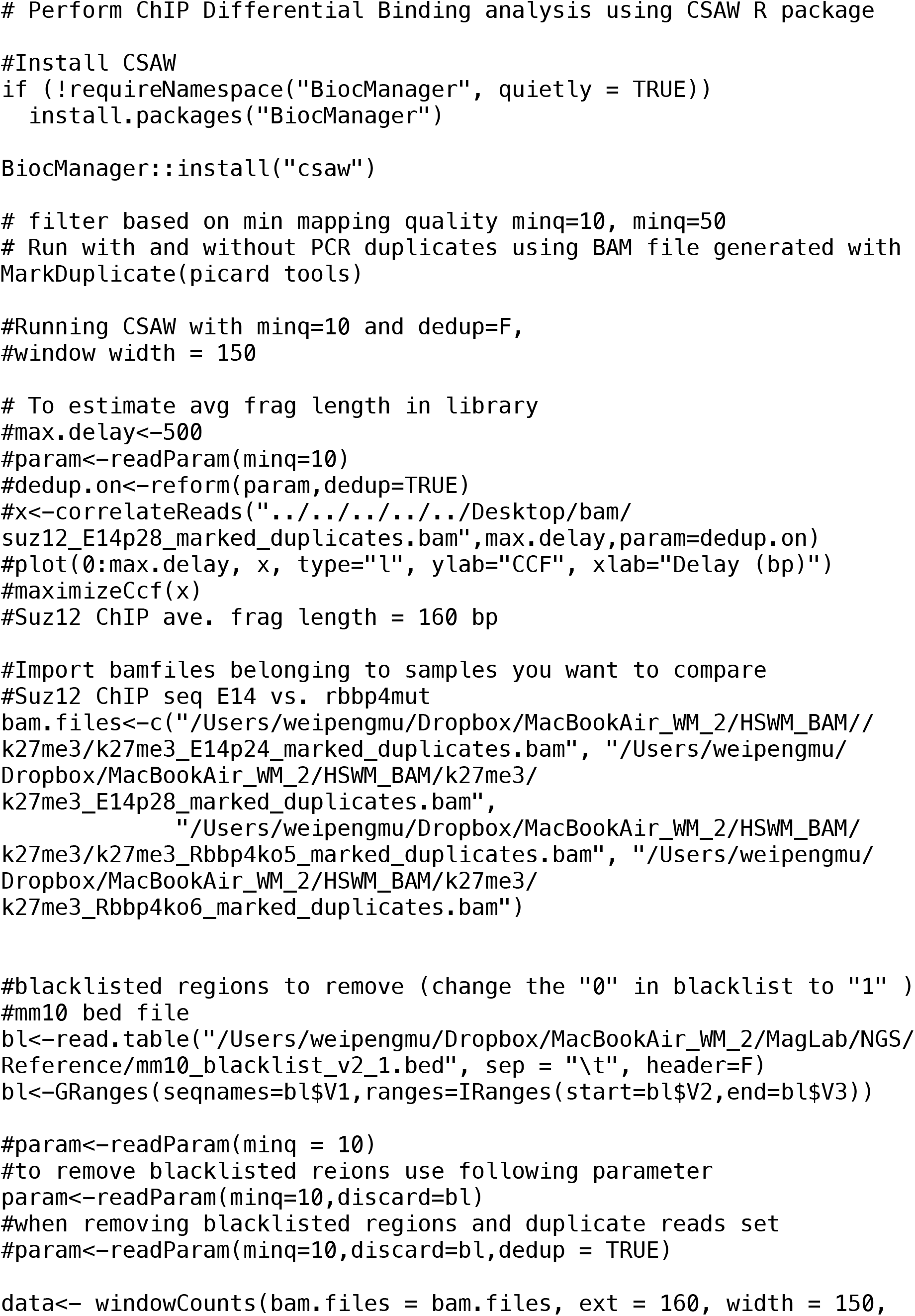
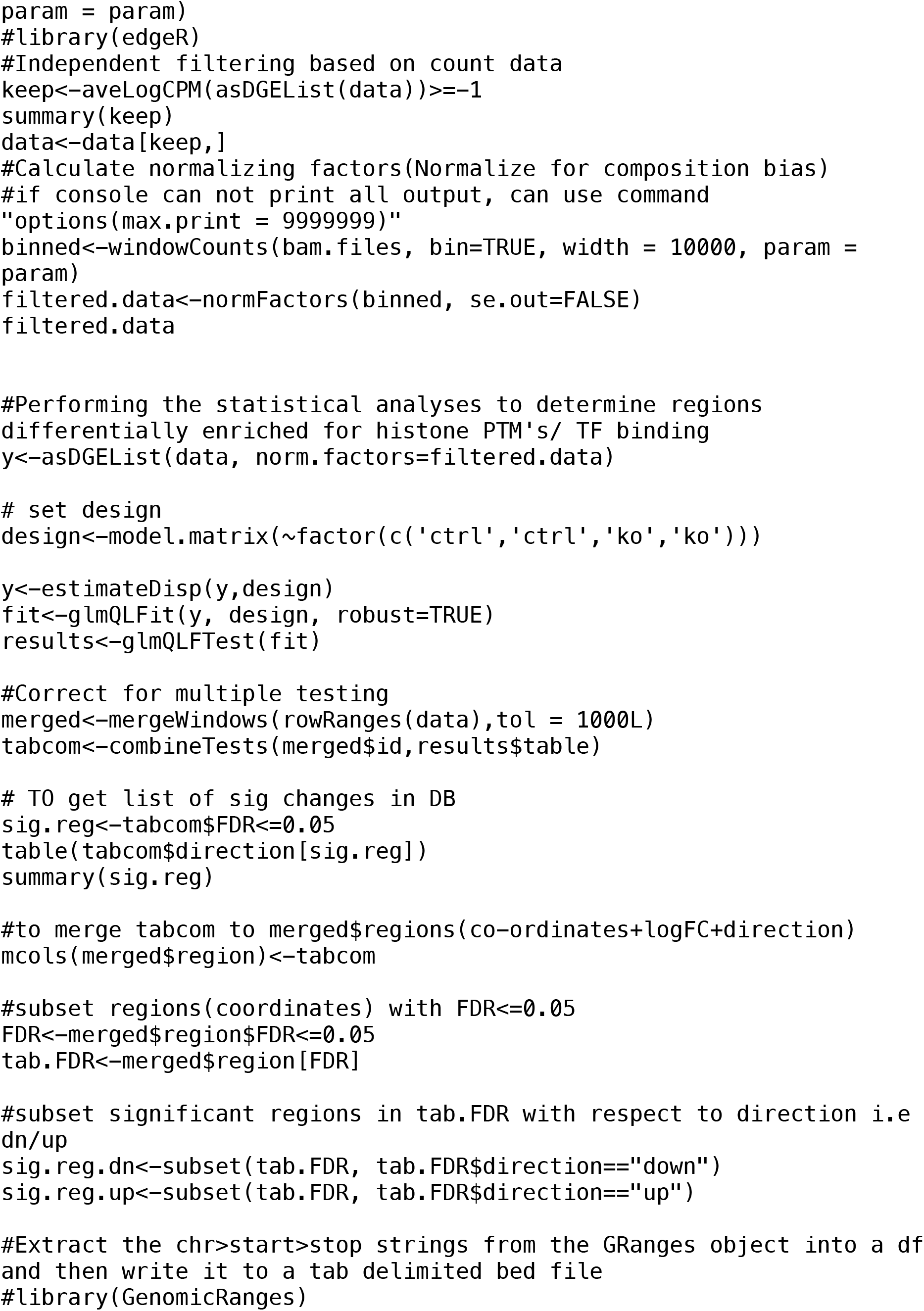
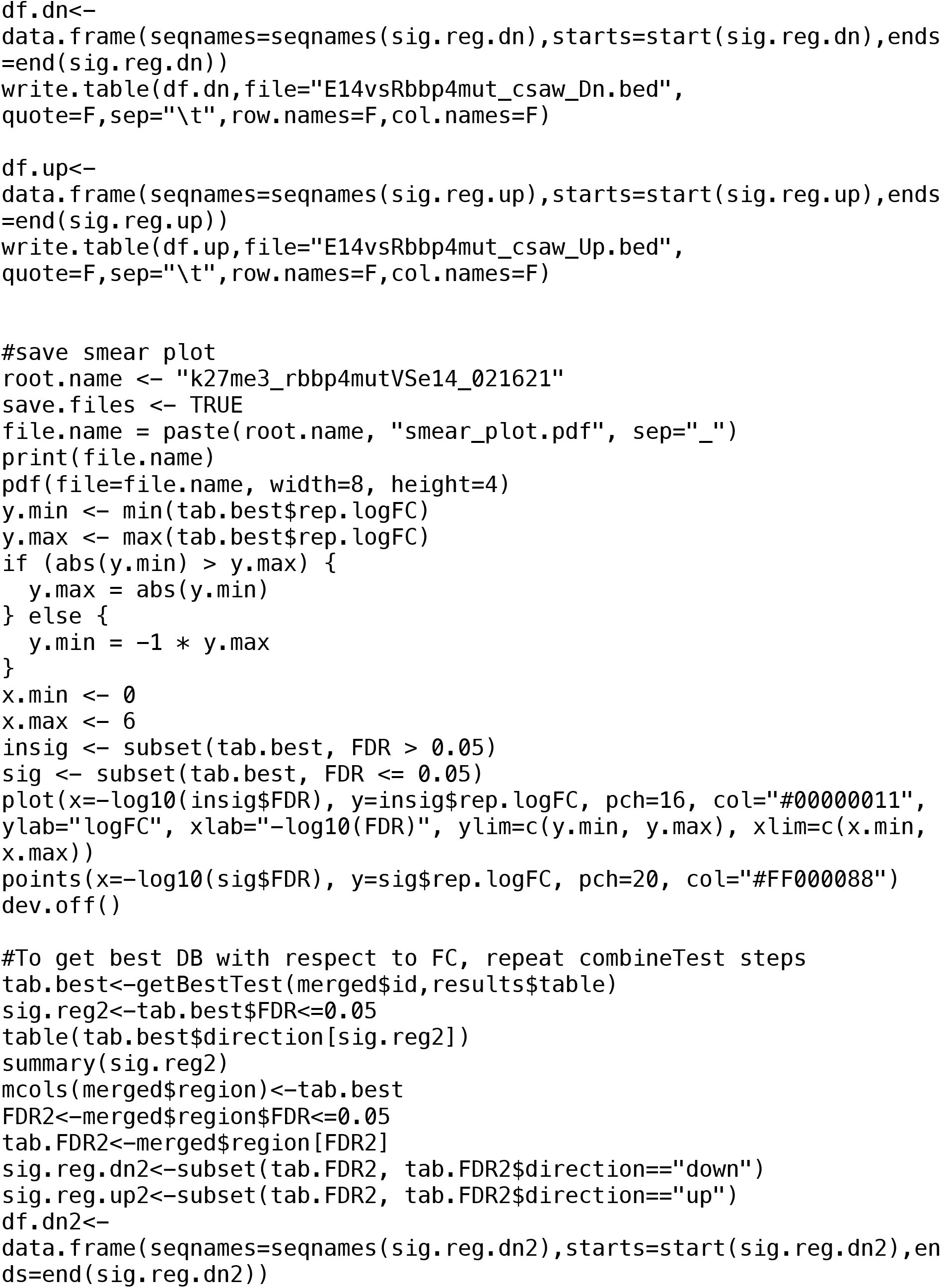
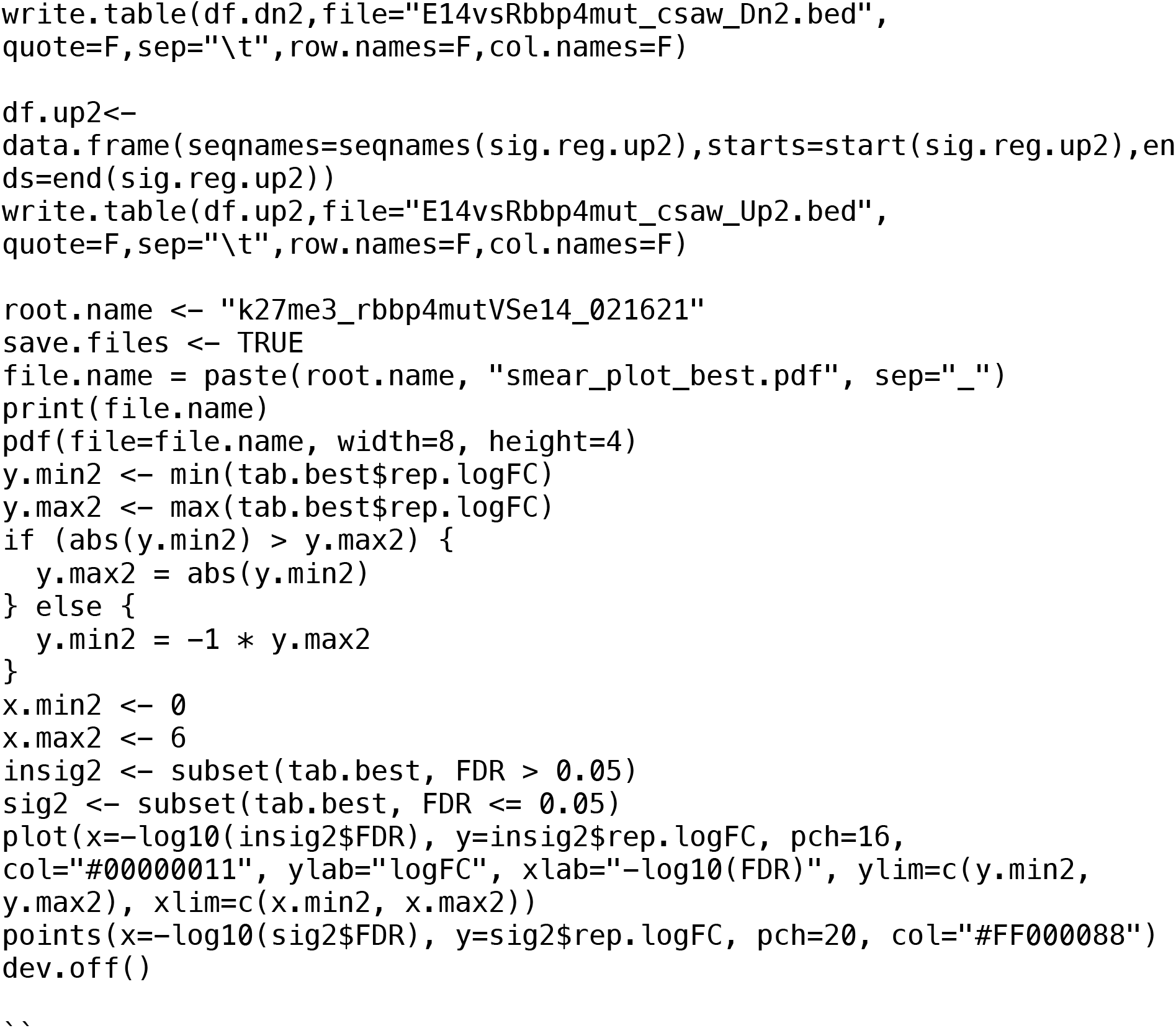

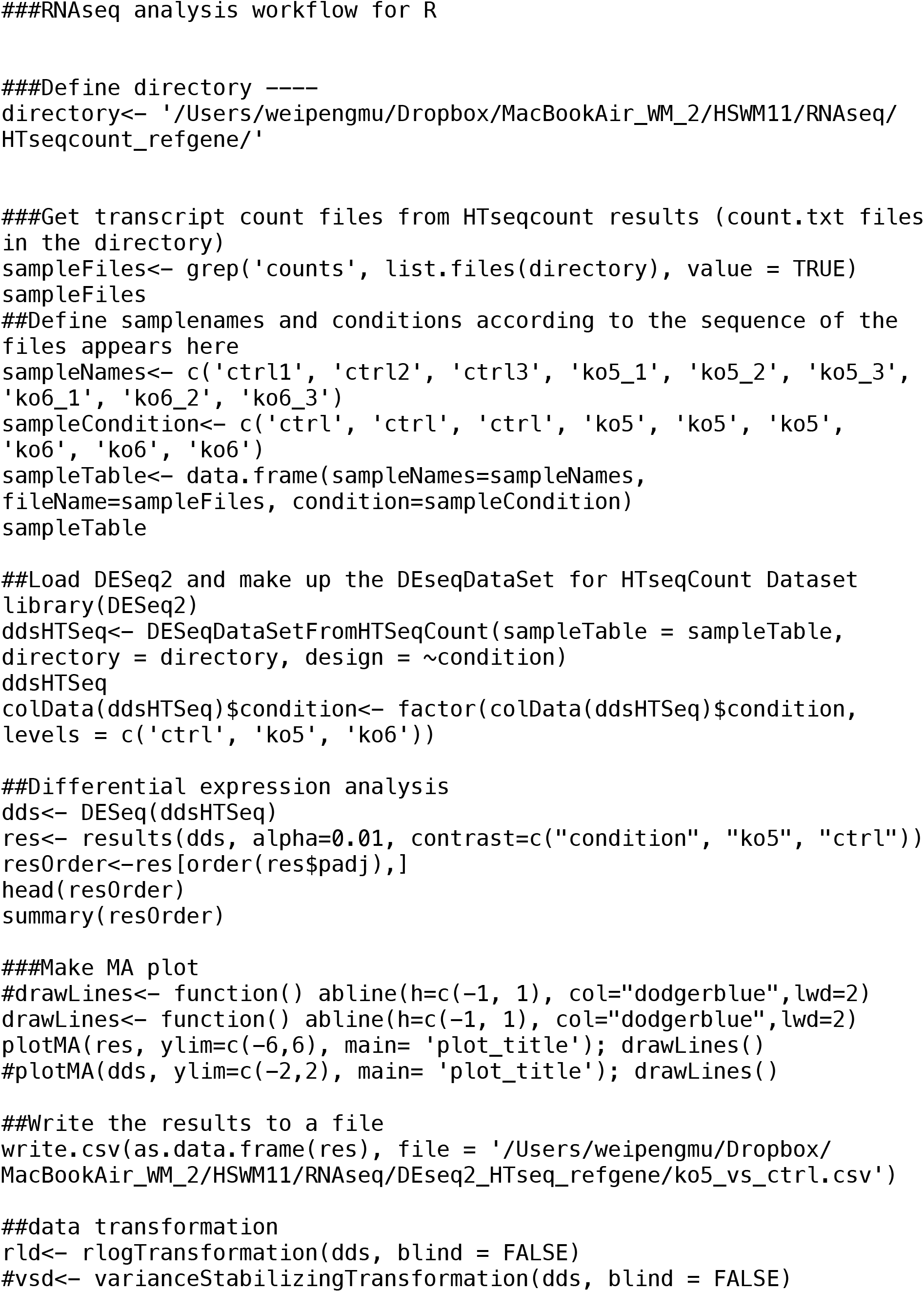
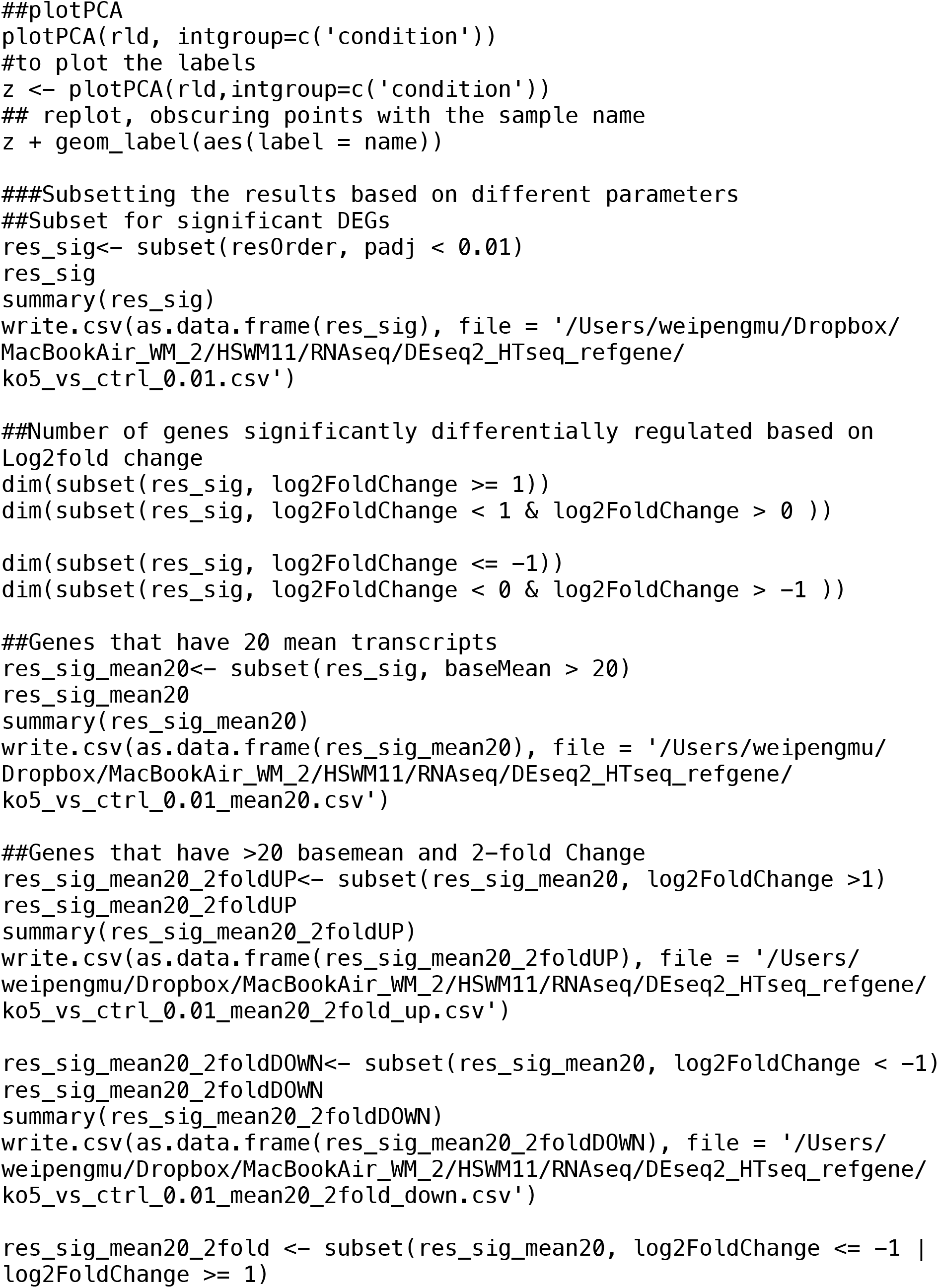
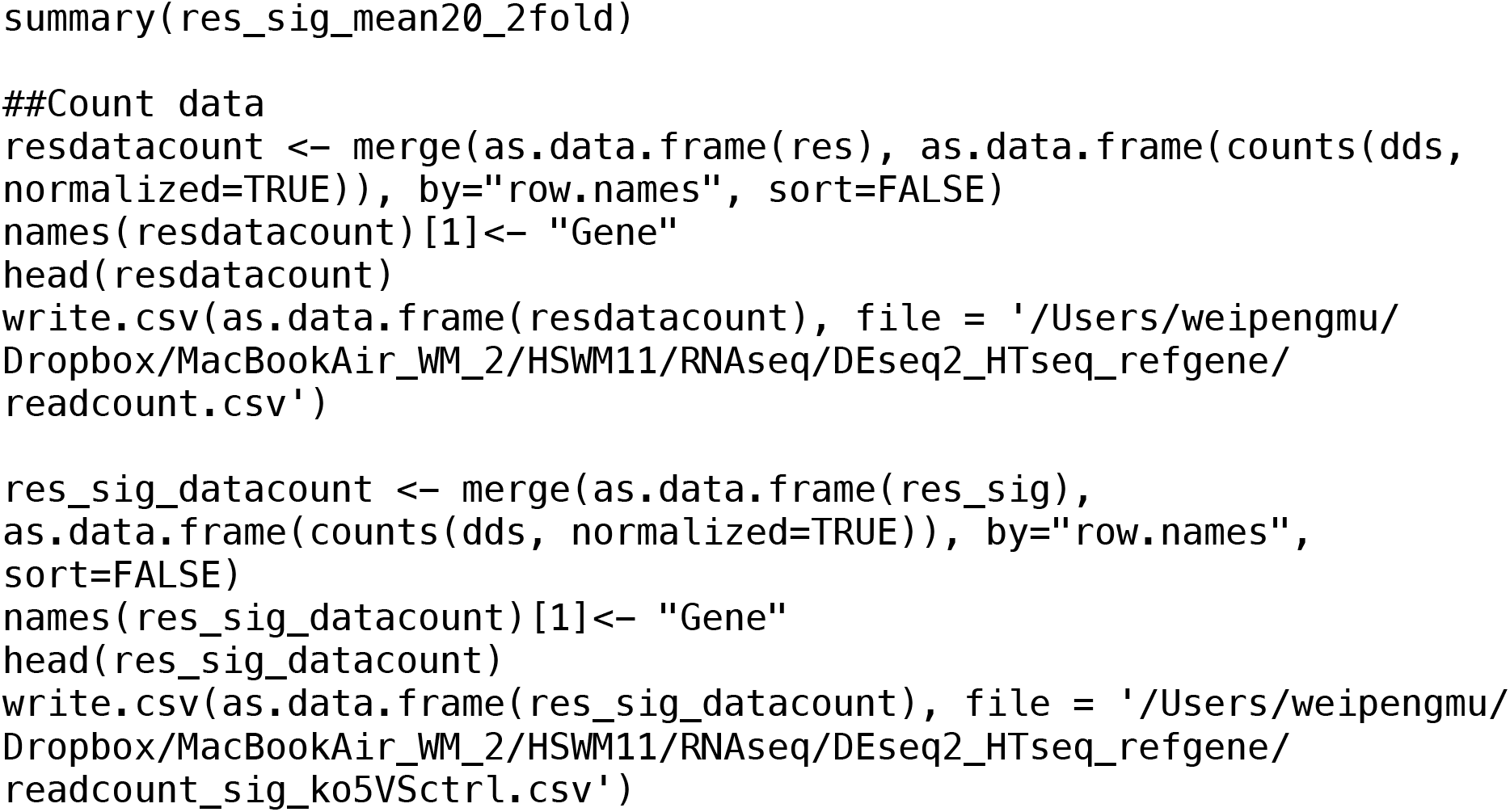

## Notes

### Competing Interest Statement

The authors have declared no competing interest.

